# Tumor Microenvironment Cellular Crosstalk Predicts Response to Adoptive TIL Therapy in Melanoma

**DOI:** 10.1101/2022.12.23.519261

**Authors:** David Barras, Eleonora Ghisoni, Johanna Chiffelle, Angela Orcurto, Julien Dagher, Noémie Fahr, Fabrizio Benedetti, Isaac Crespo, Stefan Zimmermann, Rafael Duran, Martina Imbimbo, Maria Ochoa de Olza, Blanca Navarro, Krisztian Homiscko, Sara Bobisse, Danny Labes, Zoe Tsourti, Charitini Andriakopoulou, Fernanda Herrera, Alizée Grimm, Matteo Morotti, Rémy Pétremand, Reinhard Dummer, Gregoire Berthod, Michal Bassani-Sternberg, Niklaus Schaefer, John O Prior, Maurice Matter, Nicolas Demartines, Veronica Aedo, Clarisse Dromain, Jesus Corria-Osorio, Stephanie Tissot, Lana E. Kandalaft, Raphael Gottardo, Mikael Pittet, Christine Sempoux, Olivier Michielin, Urania Dafni, Lionel Trueb, Alexandre Harari, Denarda Dangaj Laniti, George Coukos

## Abstract

Adoptive cell therapy (ACT) using *ex vivo* expanded tumor-infiltrating T lymphocytes (TILs) can mediate responses in metastatic melanoma, but long-term efficacy remains limited to a fraction of patients. Here we interrogated tumor-microenvironment (TME) cellular states and interactions of longitudinal samples from 13 metastatic melanoma patients treated with TIL-ACT in our clinical study (NCT03475134). We performed single-cell RNA-seq and spatial proteomic analyses in pre- and post-ACT tumor tissues and showed that responders exhibited higher tumor cell-intrinsic immunogenicity. Also, endogenous CD8^+^ TILs and myeloid cells of responders were characterized by increased cytotoxicity, exhaustion and costimulation and type-I IFN signaling, respectively. Cell-cell interaction prediction analyses corroborated by spatial neighborhood analyses revealed that responders have rich baseline intratumoral and stromal tumor-reactive T-cell networks with activated myeloid populations. Successful TIL-ACT therapy further reprogrammed the myeloid compartment and increased TIL-myeloid networks. Our systematic target discovery study reveals CD8^+^ T-cell network-based biomarkers that could improve patient selection and guide the design of ACT clinical trials.

**One-Sentence Summary:** Response to adoptive TIL therapy in melanoma is determined by CD8^+^ TIL-myeloid cell networks

## INTRODUCTION

Adoptive cell therapy (ACT) using *ex vivo*–expanded autologous tumor-infiltrating lymphocytes (TILs) is a potent strategy, with objective responses seen in a subset of metastatic patients with melanoma in multiple clinical studies (*1–3*) and superior to immune checkpoint inhibitor (Ipilimumab) therapy (*4, 5*). Clinical responses with TIL-ACT have also been reported in epithelial cancers (*6–8*). However, the benefit of TIL-ACT does not extend to all treated patients, for reasons that remain to date unclear.

Advances in single-cell (sc) technologies enable in-depth characterization of tumor and immune cells in the tumor microenvironment (TME) (*9*). Although the molecular profiles of CD4^+^ or CD8^+^ tumor-specific TILs at the steady state have been recently reported by single-cell (sc) RNAseq (*10–14*), those of other leukocytes regulating T-cell function or performance during ACT remain unclear. Yet, understanding the steady-state immune contexture could help improve the selection or the preparation of patients for TIL-ACT. Furthermore, no study has investigated to date, the TME dynamics during TIL-ACT in human solid tumors. Such understanding could elucidate key mechanisms supporting T-cell mediated immune rejection but also therapeutic resistance, and help elaborate future therapeutic strategies.

To identify T-cell and other TME-cell states associated with successful TIL-ACT, we carried out sc and bulk RNAseq, corroborated by multispectral immunofluorescence imaging (mIF) analysis of longitudinal samples from metastatic melanoma patients before and after TIL-ACT in a phase-I clinical study (NCT03475134). We show that response to adoptive TIL therapy in melanoma is determined by pre-existing and identifiable CD8^+^ TIL-myeloid cell networks, which support immune attack at the steady state and reorganize post TIL transfer to sustain immune rejection.

## RESULTS

## Steady state melanoma microenvironment of patients subjected to ACT

We analyzed thirteen patients with metastatic melanoma who received TIL-ACT after failing immune checkpoint blockade (ICB) therapies (**Table S1**). Patients received *ex vivo* expanded TILs (*15*) and bolus intravenous IL-2 support following non-myeloablative chemotherapy, according to established protocols (*2*) (**Fig. 1A**). Patient characteristics are detailed in Supplementary Materials. We observed objective responses in 6/13 patients (best overall response by RECIST v.1.1 at 3 months: 46%), resulting in median progression-free survival (PFS) of 5.6 months (95% CI 1.2 – 8.5), and median overall survival (OS) of 8.8 months (95% CI 7.5 – not reached, with a median follow-up of 30.2 months, IQR: 27.0 - 36.2), which compared favorably to a synthetic control arm from the same institution of clinically matched melanoma patients who received no TIL-ACT following ICB (median OS: 6.7 months) (**Fig 1B**, **Fig. S1A**). In particular, two patients attained an ongoing complete response (CR) and four a partial response (PR) at three months (classified as Responders, Rs), while three had stable disease (SD) and four progressive disease (PD) at three months (non-responders, NRs) (**Fig 1C**, **Fig. S1B**).

**Figure 1.**
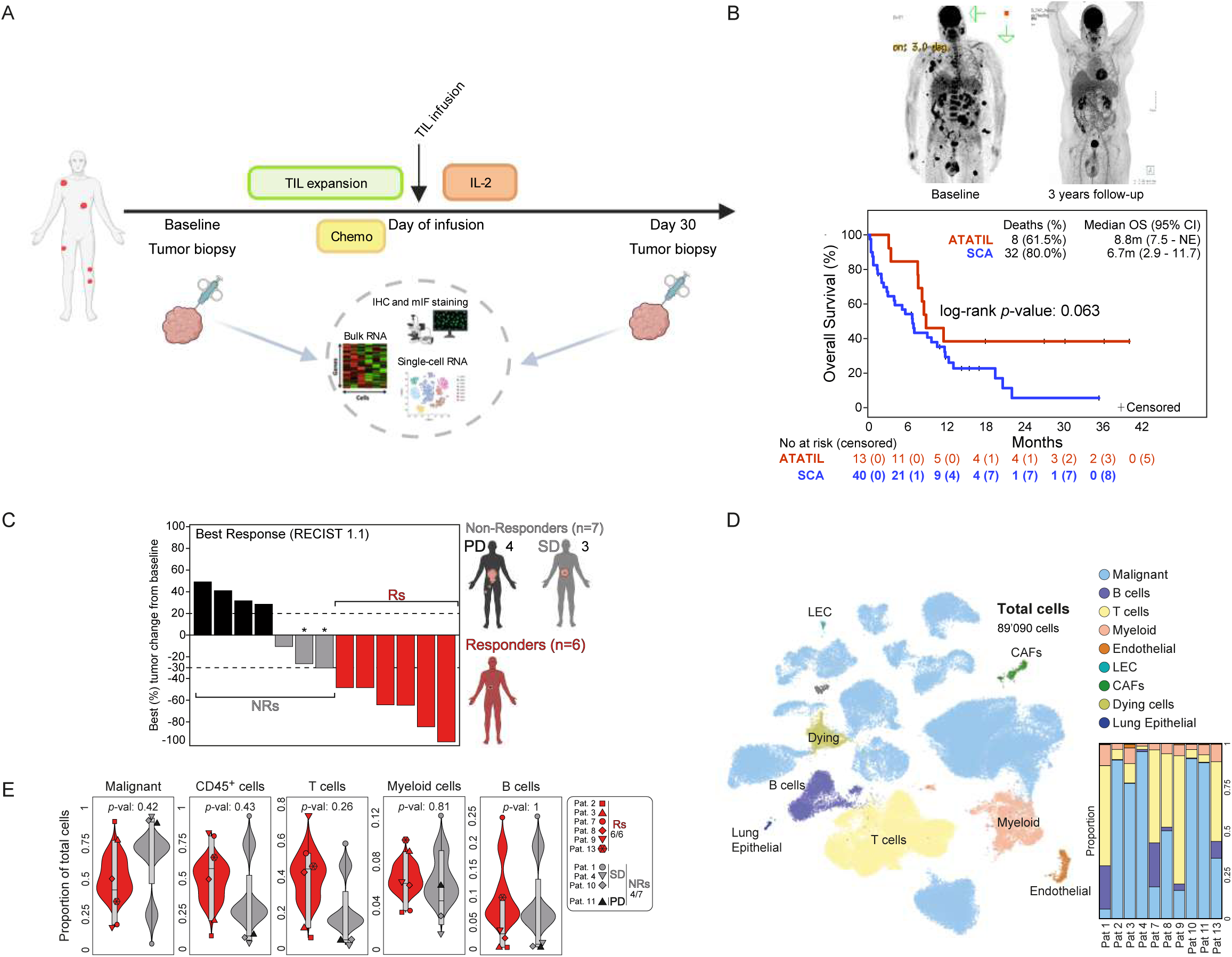
Clinical characteristics and TME landscape of our melanoma TIL-ACT cohort. (**A**) Schematic representing study design and timepoints for translational analysis. (**B)** Overall survival (OS) of our melanoma TIL-ACT cohort (NCT03475134) compared to a matched Synthetic Comparative Arm (SCA). Median follow-up time: 30.2 months (IQR: 27.0 - 36.2) by reverse Kaplan-Meyer method; log-rank *p*-value = 0.063. Top; PET-CT images of patient 3 (complete responder) at baseline and at 3 years follow-up. (**C)** Best tumor change (%) in sum of target lesions from baseline tumors. Red indicates responders (CR and PR according to RECIST 1.1 criteria), gray indicates stable disease (SD), and black indicates progressors (PD). (**D)** UMAP projection of scRNAseq data from sorted viable cells of the total TME from 10 baseline melanoma highlighting the main cell type populations and their proportions (right barplot). (**E**) Violin plots of main cell type proportions according to clinical response. Statistical significance was assessed using Student’s t-test comparing Rs versus NRs.

To understand the baseline cellular landscape of these tumors, we interrogated by scRNAseq 12 available surgical samples from 10 patients (6 Rs and 4 NRs) (**Fig. S1C**). We identified 89’090 cells distributed into 21 major clusters, including 17’965 T cells; 3’777 B cells; 4’276 myeloid cells; and other stroma or vascular endothelial cells (**Fig. 1D**, **Fig. S2A-C**). Non-malignant populations segregated in UMAP space, with cells from different patients intermixed (**Fig. S2A**). Using the Cancer Genome Atlas (TCGA) we built a melanoma-cell specific gene signature score and identified 59,958 tumor cells. These clustered separately for each patient, indicating high patient specificity **(Fig. 1D, Fig. S2A,D**). We found no differences between malignant cell or immune cell proportions based on clinical response. Even though NRs exhibited a lower trend in total TILs, this was diffused by high interpatient variability (**Fig. 1E)**.

### Baseline melanomas of responders exhibit tumor-intrinsic immunogenicity and genomic instability

We interrogated the malignant compartment at baseline (**Fig. 2A**). Copy-number variation (CNV) including amplifications and deletions, inferred from gene expression data (*16*), was significantly higher in melanoma cells from Rs relative to NRs, suggestive of higher genomic instability in Rs (**Fig. 2B-C**). These CNVs were mostly private across patients (**Fig. S2E**). Consistent with genomic instability and activation of the DNA sensing/interferon pathway (*17*), SCENIC analysis, which infers transcription factor (TF) activity based on regulon (TF and its target genes) expression (*18*), revealed several STATs and IRFs among the top activated regulons in melanoma cells of Rs (**Fig. 2D**). SOX10, activated in responders, suggested more differentiated melanoma enrichment in Rs, which was confirmed by histopathology (**Fig. S2F**). Moreover, gene set enrichment analysis (ssGSEA) showed activation of immunogenic programs in melanoma cells of Rs, including double-stranded (ds)DNA sensing, interferon (IFN)*α* and IFNγ responses, complement activation, antigen-presentation, class I/II MHC, IL-6/JAK/STAT3 signaling, TNF*α*/NF-*κ*B signaling and immune checkpoints (**Fig. 2E-F**, **Fig. S2G**). In agreement, by differential gene expression analysis we found higher expression of MHC class I/II antigen-presentation components and immunoproteasome activation in tumor cells from Rs, as evidenced by the upregulation of *B2M*, *TAP1*, *HLA-E,A,C*, *PSMB8* and *PSMB9* (**Fig. 2G**, **Table S2**). Moreover, FASL-CD95 and TRAIL apoptosis signaling pathways were overexpressed in tumor cells of responders (**Fig. 2H**, **Fig. S2H**, **Table S3**).

**Figure 2:**
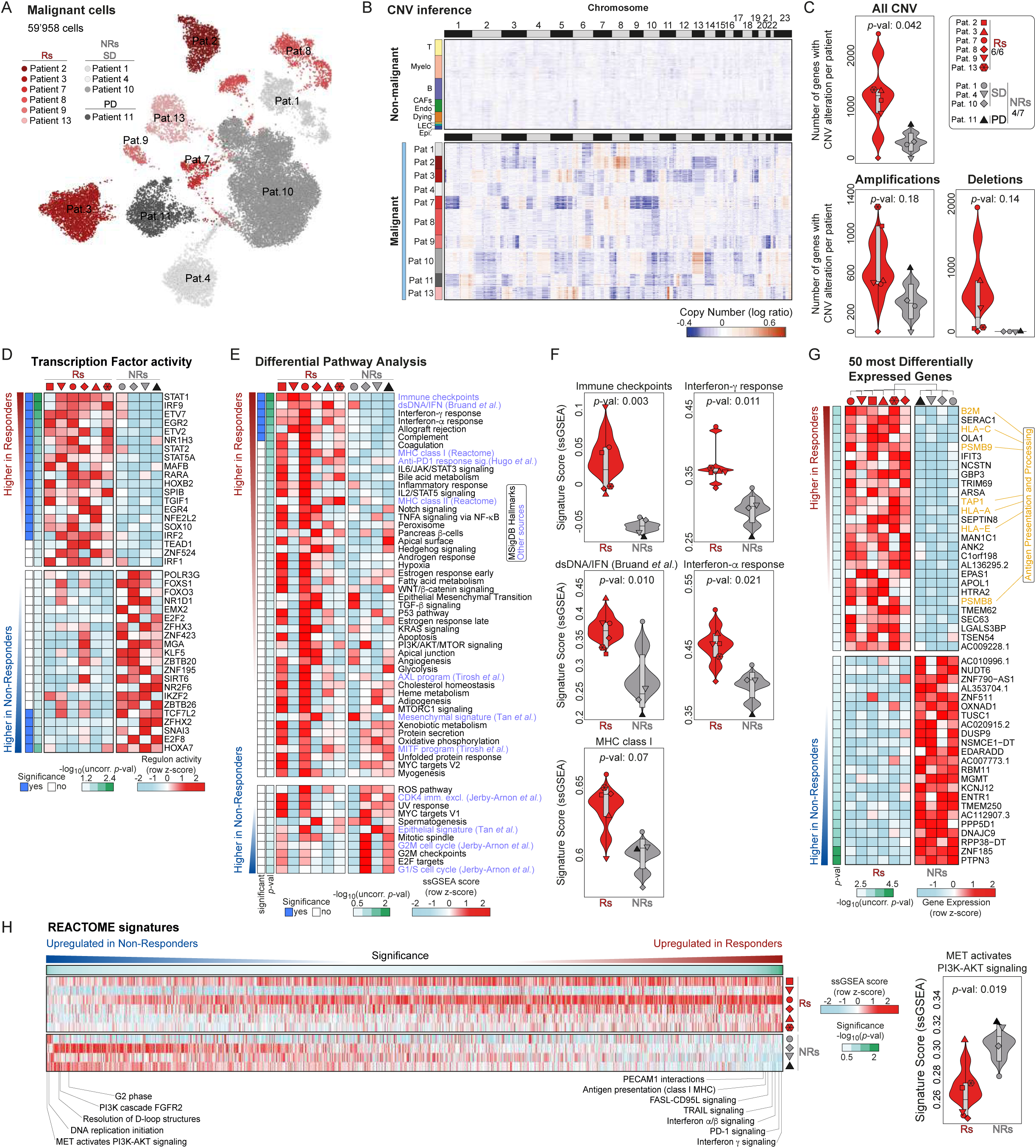
The baseline malignant compartment of TIL-ACT melanoma responders is characterized by high immunogenicity and genomic instability. (**A**) UMAP projection of malignant cells from sorted viable cell scRNAseq data of 10 baseline melanoma coloured by patient identity and clinical response. **(B**) CNV inference based on gene expression for 200 randomly selected cells from the malignant clusters of each patient. Non-malignant cells are given as reference at the top and their color label is shown in Figure 1D. (**C**) Number of genes found in (inferred) CNV-amplified and -deleted genomic regions (and the addition of both) according to clinical response. Statistical significance was assessed using Student’s t-test comparing Rs versus NRs. (**D)** Heatmap showing the most activated and repressed transcription factor/regulon in malignant cells from Rs versus NRs. Averages values of malignant cells per patient are depicted. Statistical significance was assessed using linear modeling as described in the methods. (**E)** Heatmap of Hallmarks and curated pathways that are the most differentially represented between malignant cells of Rs and NRs. Averages of malignant cells per patient are shown. Statistical significance was assessed using linear modeling as described in the methods. (**F)** Violin plots showing signature scores in malignant cells averaged per patient for the indicated Hallmarks and selected pathways according to clinical response. Statistical significance was extracted from analysis in panel E. (**G)** Heatmap showing clustering of the most up- and down-regulated significant (unadjusted *p*-value < 0.05) genes in malignant cells from Rs versus NRs. Averages of malignant cells per patient are shown. Statistical significance was assessed using linear modeling as described in the methods. (**H**) Left, Reactome pathway analysis showing selected pathways that are the most differentially represented between malignant cells from Rs and NRs. Right, Violin plots showing signature scores in malignant cells averaged per patient for the indicated Reactome pathway according to clinical response. Statistical significance was assessed using linear modeling as described in the methods.

To search for melanoma-intrinsic programs associated with lack of response to TIL-ACT, we interrogated a broader collection of Reactome signatures (**Fig. 2H**, **Fig. S2H**, **Table S3**). Among all, overexpressed in melanoma cells of NRs were the MET-driven PI3K-AKT pathway, the PI3K cascade driven by FGFR2, as well as resolution of D-loop structures, and DNA replication initiation gene expression programs, indicating DNA repair competence. Moreover, SIRT6 (NAD-dependent protein deacetylase sirtuin-6), required for genomic stability and known to control multiple glycolytic genes, was another hyperactive regulon in baseline tumors of NRs **(Fig. 2D**).

### Melanomas responding to TIL-ACT exhibit pre-existing CD8^+^ T-cell immunity

We next examined the T-cell compartment in these tumors at baseline. The proportion of overall CD4^+^ and CD8^+^ TILs or CD45^+^ leukocytes was not significantly different between Rs and NRs (**Fig. S3A**, **Fig. 1E**). By scRNAseq performed on CD45^+^-sorted cells from all 13 patients, we identified nine different CD8^+^ TIL clusters, which we assigned based on prior knowledge (*10, 19–22*) to known states (**Fig. 3A-B**, **Fig. S3B-C**, **Table S4**), including naive-like; effector-memory (EM)-like; *CX3CR1*^+^; heat-shock gene-positive (HSP); *FOXP3*^+^; interferon-stimulated gene (ISG) positive; precursor exhausted (Pex); and exhausted (Tex). Additionally, we identified NK-like CD8^+^ TILs; a NK-cell cluster; and three CD4^+^ T-cell subsets: T-helper 1 (Th1); *CXCL13*^+^ T-follicular helper (Tfh)-like; and T-regulatory (Treg) cells (**Fig. S3D-E**, **Table S4**). State phenotypic divergence of CD8^+^ TILs inferred by pseudotime differentiation trajectories suggested that *CX3CR1*^+^ and exhausted (and a fraction of ISG and EM-like) CD8^+^ TILs were the most differentiated effector states, but represented distinct programs (**Fig. 3C**, **Fig. S3F**). Together, these three states contributed equally to collectively one-third of all CD8^+^ TILs recovered, while EM-like cells contributed another third.

**Figure 3:**
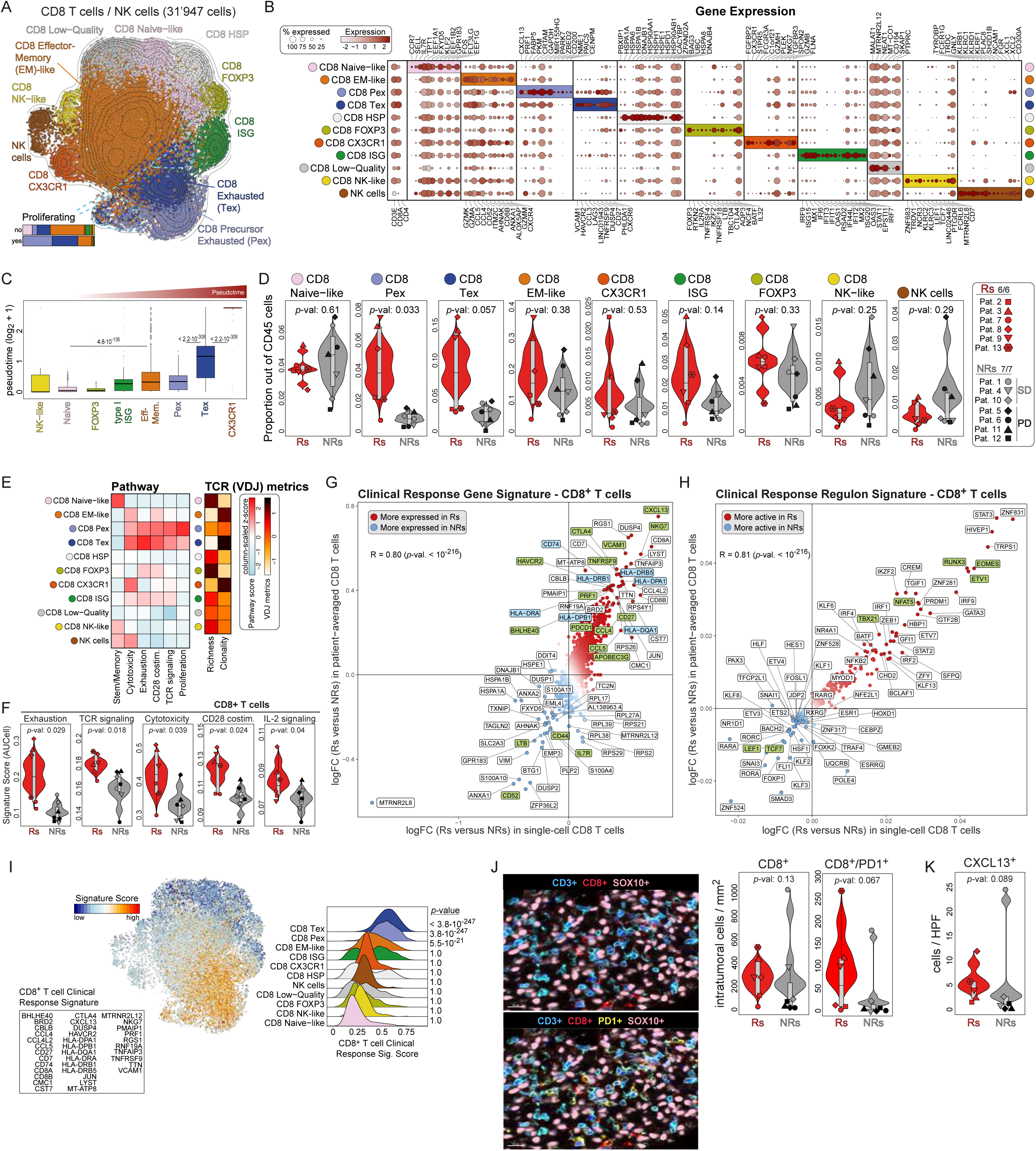
CD8^+^ T cells of ACT responders exhibit higher cytotoxicity, exhaustion and costimulation transcriptional programs at baseline. (**A**) UMAP projection showing sub-clustering of CD8^+^ T and NK cells from CD45^+^-cell sorted data of thirteen baseline and seven post-ACT tumors. (**B**) Characterization of the CD8^+^ T cell populations; dot plot showing gene expression for the most discriminant gene markers. (**C**) Pseudotime analysis of the CD8^+^ T cell populations showing pseudotime values per cell grouped per CD8^+^ T cell subtype. Statistics performed using ANOVA followed by *post-hoc* Tukey test. (**D**) Violin plots of CD8^+^ T cell subtypes and NK cell proportions over CD45^+^ cells in CD45^+^-cell sorted data according to clinical response. Statistical significance was assessed using Student’s t-test comparing Rs versus NRs. (**E**) Left: gene signature scores for stemness/memory, cytotoxicity and exhaustion (taken from Jerby-Arnon *et al*.), TCR signaling and costimulation mediated by CD28 (Reactome) and proliferation per CD8^+^ T cell subtype. Right: richness and clonality TCR (VDJ) metrics as computed using scTCRseq data. **(F**) Violin plots showing cytotoxicity and exhaustion (taken from Jerby-Arnon *et al*.), TCR signaling, IL-2 signaling and costimulation mediated by CD28 (Reactome) scores for CD8^+^ T cells of the CD45^+^-cell sorted data according to clinical response. Statistical significance was assessed using Student’s t-test comparing Rs versus NRs. (**G-H)** Differential gene expression (**G**) and transcription factor/regulon (**H**) analysis between CD8^+^ T cells from Rs versus NRs. The x-axis displays the log2 fold-change as computed at single-cell level while the y-axis shows the log2 fold-change as computed in patient-averaged data. Statistical analysis was assessed using linear modeling as described in the methods. (**I**) Left: box displaying the CD8^+^ T cell clinical response gene signature that was obtained by the union of the 30 most upregulated genes in Rs at single-cell and patient-averaged level (from panel G). Scoring of the CD8^+^ T cell clinical response signature projected in UMAP space (middle) and plotted as ridge plot per CD8^+^ T cell subsets. Statistics performed using one-sided Student’s t-tests of each population versus all the others and after Bonferroni correction. (**J**) Example and quantification of multi-immunofluorescence imaging (mIF) staining showing intratumoral (SOX10^+^), CD3^+^, CD8^+^, PD-1^+^ cells at 10x magnification (left). Violin plots of cell density (cell/mm^2^) between Rs and NRs (right). Statistics performed using one-sided wilcoxon tests (right). (**K**) Violin plot of CXCL13^+^ cell density (cell count/HPF) between Rs and NRs (at 20x magnification). Statistics performed using one-sided wilcoxon tests.

Recently, human tumor-antigen specific CD8^+^ TILs were found distributed across three *TOX*^+^*PDCD1*^+^*CTLA4*^+^*TIGIT*^+^ transcriptional exhaustion states, including a precursor state (high expression of *TCF7*, *LEF1*, *CCR7* and *IL7R*); a terminally differentitated state (high *HAVCR2*/TIM3 and *ENTPD1*/CD39 and cytolytic effector molecules), and an intermediate state (*23*) defined by TFs of recent TCR engagement. Here, we found a significant enrichment in CD8^+^ TILs at similar Pex and Tex states in Rs, along with (non-significantly) higher frequencies of ISG CD8^+^ TILs (**Fig. 3D**). Consistent with the pseudotime-based differentiation inference, Tex, EM-like and *CX3CR1*^+^ cells exhibited by scTCRseq the most oligoclonal expansion, suggesting antigen selection or tumor specificity (*24–26*). However, among all cells, Pex and Tex exhibited – in addition to exhaustion – the highest signatures of TCR signaling and cytotoxicity as well as proliferation (**Fig. 3E**), indicating tumor antigen engagement.

Consistent with a precursor state and increased proliferative capacity, Pex TILs overexpressed the most DNA amplification and DNA repair pathways as well as proliferation signatures (**Fig. 3E**, **Fig. S3G**), and relative to Tex displayed higher inferred regulon activity of TCF7, MYC, MYOD1, and HDAC2 chromatin modulators, while conversely Tex exhibited higher RUNX3 (*27*) regulon activity, a key regulator of CD8^+^ T-cell tissue residence (*28*) and Tex effector function during chronic viral infection (**Fig. S3H**). Consistent with previous findings in ovarian cancer that a fraction of exhausted CD8^+^ TILs retain proliferative capacity and polyfunctionality (*29*) as a consequence of CD28 costimulation received by DCs *in situ*, we found that also melanoma Tex (and Pex) exhibited signatures of CD28 costimulation and proliferation in Rs at baseline (**Fig. 3E**). Interestingly, exhausted CD8^+^ TILs with lower levels of CD28 costimulation exhibited lower EOMES and TBX21 activity (**Fig. S3I**), implying that a subset of Tex cells may not have transitioned to terminal exhaustion (*30, 31*), as suggestd also by proliferation. Indeed, TILs transition through a continuum gradient of exhaustion states (*26*) as also implied by our pseudotime analysis (**Fig. S3F**).

ISG CD8^+^ TILs, distinguished by a dominant signature of IFN response genes, have been previously reported in human TILs (*25*). Here they exhibited overlap in oligoclonally expanded TCR sequences with exhausted CD8^+^ TILs (**Fig. S3J**), and displayed gene signatures of cytotoxicity, TCR signaling and exhaustion, implying that ISG cells are antigen-experienced tumor-reactive Tex, located in IFN-rich tumor niches.

Strikingly, a fraction of EM-like CD8^+^ TILs exhibited the highest level of clonal TCR overlap with the exhausted compartment, indicating that these cells are also tumor-specific, but likely not yet engaged in antigen recognition as they lacked signatures of TCR signaling or exhaustion. Indeed, human *GZMK*^+^ EM TILs can transition into the Tex compartment (*25*). Remarkably, the frequencies of EM-like cells were similar in Rs and NRs at baseline (**Fig. 3D**), likely reflecting the fact that an important proportion of these cells are made of unrelated bystander cells. Finally, consistent with *CX3CR1*^+^ TILs not contributing significantly to antitumor immunity in the mouse (*32*), here we found that *CX3CR1*^+^ CD8^+^ TILs exhibited no clonotype overlap with the exhausted compartment nor TCR signaling or exhaustion programs, indicating no role in tumor recognition. Indeed, close interrogation of the gene profile and inferred regulon activity in these cells revealed that they are not the human equivalent to the mouse intermediate exhausted CX3CR1^+^ cells with cytolytic potential, which are also PD-1^int^, TOX^int^ and T-bet^hi^ (*28, 33*). Since we found that the human *CX3CR1*^+^ cells were closer to EM based on trajectory analysis, we infer that these are most likely bystander EM cells (*25*). Indeed, their frequency was similar in Rs and NRs (**Fig. 3D**, **Fig. S3F)**.

Matching the above findings, baseline CD8^+^ TILs were enriched for transcriptomic programs of exhaustion, TCR signaling, cytotoxicity, CD28 costimulation, IL-2 signaling, IFNγ activation and effector function (**Fig. 3F**, **Fig. S3K**). To learn more, we analyzed differentially expressed genes in CD8^+^ TILs from Rs and NRs, both at a single-cell and at a patient-averaged level. Their correlation revealed highly consistent overexpressed genes (*CXCL13*, *CCL5*, *HLA-DRB5*, *HAVCR2*, *TNFRSF9*, *CTLA4*, *PDCD1*) and regulons (RUNX3, EOMES, TBX21, NFAT5, ETV1) in CD8^+^ TILs of responders (**Fig. 3G-H**, **Table S2**), many of which were previously associated with exhausted/CD28-costimulated CD8^+^ TIL signatures (*29*). Moreover, clinical response was associated with baseline CD8^+^ TILs that exhibited increased inferred activity of ZNF831 (zinc finger protein 831) and HIVEP1 (zinc finger protein 40), which bind to the enhancer elements of promoters of several key genes implicated in the effector and exhaustion programs, including the IL-2 receptor and IFNβ; *TBX21*/T-bet; *EOMES*; *PRDM1*/BLIMP-1 implicated in transcriptional activation of multiple immune checkpoint genes; *ETV1* which targets TOX (33); and *RUNX3* (**Fig. 3H**). Conversely, failure of TIL-ACT was associated with predominance of naive and memory-like CD8^+^ TILs at baseline, highlighted by the overexpression of *IL7R* and *LTB* as well as higher inferred activity of different TFs implicated in the WNT/β-catenin or TGF-β signaling pathways including LEF1, TCF7, FOXP1 and SMAD3, respectively (**Fig. 3G-H**, **Table S2**), reflecting the paucity of antigen-experienced tumor-reactive populations. Finally, we computed a CD8^+^ T-cell signature score associated with clinical response from **Fig. 3G** (**Table S2**). CD8^+^ Tex and Pex, followed by EM-like, were the TIL subsets that most highly overexpressed the T-cell clinical response signature (**Fig. 3I)**, confirming their relevance for TIL-ACT.

We corroborated these findings by multispectral immunofluorescence (mIF) microscopy and immunohistochemistry (IHC) interrogation of baseline tumors, showing higher frequencies of intratumoral CD3^+^CD8^+^ TILs (located within tumor nests) expressing PD-1 or CXCL13 in Rs (**Fig. 3J-K**, **Fig. S3L-M**). Thus, *in situ* T-cell activation, cytotoxicity, costimulation and exhaustion along with immunogenic tumor cells are hallmarks of response to TIL-ACT.

### Other baseline lymphoid states in responders and non-responders

Tfh-like CD4^+^ *CXCL13*^+^ and Treg cells were also higher at baseline in Rs (albeit not significant), completing an immune contexture of activated T-cell immunity (**Fig. S3N**). Interestingly, we identified highly proliferative γδ T cells expressing *TRGC2* and *TRDV1/TRDC* TCR chains, KLR molecules (*KLR D1*/*K1*/*C2*/*C4*/*G1*), *CCL5, NKG7* and granzymes (**Fig. S3E,O**), which were significantly enriched in NR tumors, suggesting a negative influence on TIL-ACT (*34, 35*) (**Fig. S3P).**

The important role of tumor-resident B cells in supporting antitumor T-cell responses and response to ICB is being increasingly recognized. In addition to plasma cells (*MZB1*^+^), we identified follicular-like (*MS4A1*^+^) tumor-resident CD20^+^ B cells clustering in three main states: naive (*TCL1A*^+^, *FCER2*^+^, *IGHD*^+^); memory (*CD27*^+^); and germinal-center (GC) (CD38^+^, *MEF2B*^+^) (*29*) (**Fig. S4A-B**, **Table S4)**. Memory B cells displayed high levels of IFN signaling and mature-APC genes (*CD80*, *CD86* and *CD40*), while GC B cells overexpressed signatures of B-cell receptor (BCR) signaling, class-II antigen presentation and proliferation (**Fig. S4B)**. Although we could not detect significant differences in the proportions of individual B-cell subsets between Rs and NRs **(Fig. S4C),** collectively B cells from Rs, particularly memory cells, displayed increased interferon signaling and MHC class-II antigen presentation, implying a possible participation in modulating local T cell immunity (**Fig. S4A-B, Table S4**).

### TIL-ACT responding melanoma are infiltrated by activated macrophages and dendritic cells

A mounting body of evidence indicates that the organization of the tumor myeloid compartment is crucial for orchestrating T-cell immunity. For example, tumor-resident myleoid cells promote intratumoral T-cell engraftment (*36*) and support CD8^+^ Tex effector functions via CD28 costimulation (*29*), while macrophage states predict response to checkpoint immunotherapy (*37–39*). On the basis of scRNAseq data obtained from all patients, we detected 11 transcriptional clusters of myeloid cells, which we annotated considering well-described datasets (*40–42*) into seven monocyte/macrophage states (*CXCL9*^+^, ISG, *S100A8*^+^, *TREM2*^+^ and *C1Q^+^* macrophages, monocytes, and monocyte-like-DCs (monoDC) and four dendritic (DC) states (DC1, DC2, *CCR7^+^* DC3 and plasmacytoid (p)DCs). Each of these states was also characterized by distinct inferred regulon activities (**Fig. 4A**, **Fig. S4D-E**, **Table S4**).

**Figure 4:**
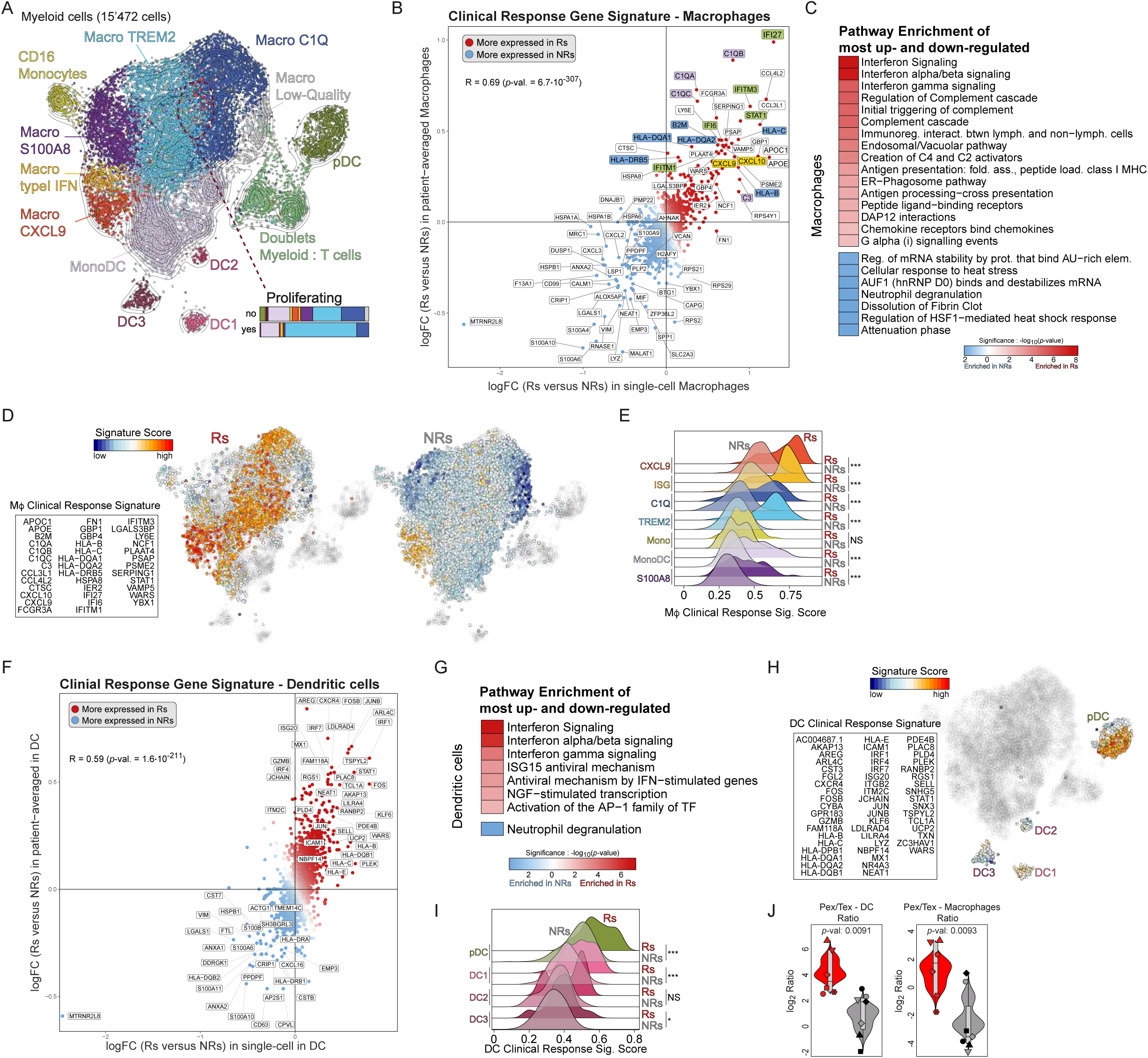
Complement activation, interferon-inducible chemokines, antigen presentation and CD28 costimulation, dominate the phenotype of macrophages and DCs associated with response to TIL-ACT. (**A**) UMAP projection showing sub-clustering of myeloid cells from CD45^+^-cell sorted data from thirteen baseline and seven post-ACT tumors. (**B**) Differential gene expression analysis between macrophages from Rs versus NRs. The x-axis displays the log fold-change as computed at single-cell level while the y-axis shows the log fold-change as computed in patient-averaged data. Statistical analysis was assessed using linear modeling as described in the methods. (**C**) The 30 most upregulated and downregulated genes in macrophages between Rs and NRs (from Panel B) were subjected to Reactome enrichment analysis and significant (false discovery rate < 0.01) pathways appear for the genes upregulated in responders (in red) or upregulated in the non-responders (in blue). (**D**) Left: box displaying the macrophage clinical response gene signature that was obtained by the union of the 30 most upregulated genes in Rs at single-cell and patient-averaged level (from panel B). Scoring of the macrophage clinical response signature projected in UMAP space (right, split for Rs and NRs). (**E**) Ridge plot displaying the macrophage clinical response signature score per macrophage subsets split according to response. Statistics performed using Bonferroni corrected Student’s t-tests. (**F**) Differential gene expression analysis between dendritic cells from Rs versus NRs. The x-axis displays the log fold-change as computed at single-cell level while the y-axis shows the log fold-change as computed in patient-averaged data. Statistical analysis was assessed using linear modeling as described in the methods. (**G**) The 30 most upregulated and downregulated genes in dendritic cells between Rs and NRs (from panel F) were subjected to Reactome enrichment analysis and significant (false discovery rate < 0.01) pathways are shown for the genes upregulated in responders (in red) or upregulated in the non-responders (in blue). (**H**) Left: box displaying the DC clinical response gene signature that was obtained by the union of the 30 most upregulated genes in Rs at single-cell and patient-averaged level (from panel F). Right: scoring of the DC clinical response signature projected in UMAP space. (**I**) Ridge plot displaying the DC clinical response signature score per DC subsets split according to clinical response. Statistics performed using Bonferroni corrected Student’s t-tests. (**J**) Log_2_ ratio of the proportions between exhausted T cells (union of CD8 Pex and Tex) and bulked DC or macrophage populations in single-cell data of baseline tumors. Initial proportions were taken out of CD45^+^ cells. Statistical significance was assessed using Student’s t-test comparing Rs versus NRs.

When comparing myeloid cells from Rs and NRs, we observed no significant differences in the relative proportions of DCs but an important enrichment for monocytes/macrophages among leukocytes in NRs (**Fig. S4F**). Also, we observed no statistically significant differences when analyzing each of the 11 individual myeloid cell states (**Fig. S4F**). Considering that macrophage polarization may have either anti- or pro-tumor effects (*39*), we investigated whether such polarization differed between Rs and NRs. Because macrophages can express M1- and M2-like phenotypes (**Fig. S4E**) that have been associated with their ability to promote or suppress T cells respectively, at least *in vitro*, we asked whether the ratio of canonical M1 and M2 markers in macrophages at baseline was associated with clinical response. Using M1 and M2 gene signatures (*43*), we assigned each macrophage-cell to an M1 and/or M2 phenotype and quantified their ratios within the macrophage compartment (**Fig. S4G**). We found that Rs had a higher ratio of M1/M2 compared to NRs (**Fig. S4G**).

To learn more, we analyzed differentially expressed genes in macrophages from Rs and NRs, both at a single-cell and at a patient-averaged level. Their correlation unveiled robustly overexpressed genes (**Fig. 4B**), specifically revealing in responders’ macrophages interferon signaling (*IFI27*, *IFITM1, IFITM3*, *IFI6, STAT1*), upregulation of IFN-inducible chemokines *CXCL9* and *CXCL10*, complement component synthesis (*C1QA-C*, *C3*), phagocytosis, and antigen processing and presentation (*HLA-B*, *HLA-C*, *B2M*, *HLA-DQA1*, *HLA-DQA2*, *HLA-DRB5*) (**Fig. 4B-C**, **Table S2-3**). We next computed a clinical response-associated macrophage score using the most upregulated genes in Rs (**Fig. 4D**, **Table S2**) and interrogated the different macrophage subsets. Revealing their relevance for TIL-ACT, this signature was highly overexpressed in *CXCL9*^+^ and ISG macrophages followed by *C1Q*^+^ and *TREM2*^+^ cells, specifically in Rs (**Fig. 4E**, **Fig. S4H**). CXCL9 in macrophages is specifically upregulated by IFNγ (*36*), revealing the presence of nearby TILs recognizing tumor antigen, while it further amplifies intratumoral TIL engraftment (*36*). Since both *CXCL9*^+^ and ISG macrophages overexpress *CXCL9*, they are likely engaged in important crosstalks with tumor-specific T cells *in situ* at baseline. As expected, *S100A8*^+^ macrophages, which have been associated with tumor promotion and express immunosuppressive phenotypes (**Fig. S4D-E**), did not contribute to the gene signature associated with clinical response (**Fig. 4D-E**).

Next, we conducted similar analyses for DCs. As for macrophages, DCs exhibited increased activation and interferon response signatures (*IRF4-7*, *CXCR4*, *ICAM1*, *STAT1*) in Rs (**Fig. 4F-G, S4D-E**, **Table S2**), revealing an overall type-1 polarization of the myeloid compartment. We next derived an integrated DC-specific clinical response signature as above. Strikingly, this was largely confined to pDCs in Rs (**Fig. 4F,I**). While the role of pDCs in cancer is not clear presently, the presence of pDCs in human colon cancer was recently associated with increased progression-free and overall survival (*44*).

Finally, we examined the relative abundance of tumor-reactive CD8^+^ TILs over myeloid subsets, focusing on exhausted TILs (Tex-Pex combined). The frequency of exhausted TILs relative to macrophages or DCs was significantly higher in Rs, while a higher abundance of myeloid populations over T cells was associated to lack of response to ACT (**Fig. 4J**). In conclusion specific transcriptional programs, such as complement activation, interferon signaling, IFN-inducible chemokines, antigen presentation and CD28 costimulation, dominate the phenotype of macrophages and DCs which are originally enriched in tumor-reactive TILs.

### Melanomas responding to TIL-ACT are uniquely characterized by a rich immune cell crosstalk

The above findings raise the possibility that important cellular crosstalks established between TILs and tumor APCs at baseline provide the bases for successful TIL-ACT. To learn more, we inferred cell-to-cell crosstalks through ligandome analysis, computing receptor-ligand gene expression and relative proportions of putative cell pairs in the TME. These analyses revealed far more immune cell-to-cell interactions in tumors of Rs relative to NRs at baseline (**Fig. 5A**). Strikingly, in Rs we identified a plethora of putative interactions implicating CD8^+^ and CD4^+^ TIL subsets with myeloid subsets (**Fig. S5A**, **Table S5**). The only few interactions identified in NRs were between myeloid cells and tumor cells, specifically involving *S100A8*^+^ and *TREM2*^+^ macrophages, i.e. the two macrophage subsets identified as expressing uniquely M2 but no M1 signatures (**Fig. S4E**), in addition to transitory monoDCs and DC2s, but TILs appeared not involved.

**Figure 5:**
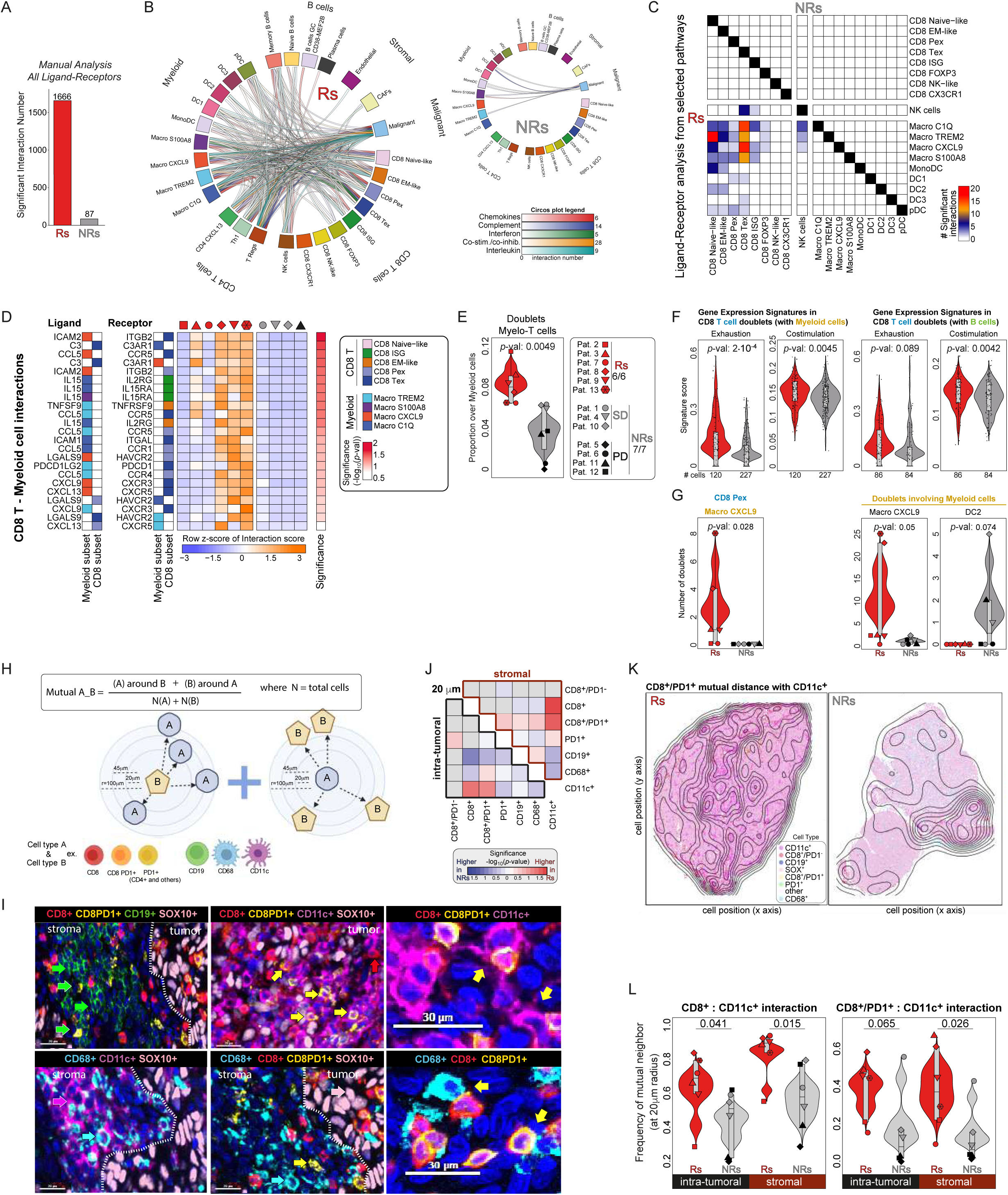
Baseline tumors of TIL-ACT responders are characterized by high T cell-TME interaction in contrast to melanoma of non-responders. (**A**) Number of total inferred interactions detected in the CD45^+^-cell sorted data using our own methodology (see Methods) according to clinical response. (**B**) Circos plots showing the finer cell subtypes involved in significant interactions (link) and the number of significant interactions (in color intensity) for the ligand-receptor pairs of five different pathways and split by clinical response at baseline. (**C**) Heatmap displaying the number of significant ligand-receptor (from selected pathways like in panels B to D) interactions between cell subsets in responders (lower-left part of the heatmap) and non-responders (higher-right part of the heatmap). (**D**) Heatmap displaying selected significant pairs of ligand-receptor (related to panel C), the myeloid and CD8 T cell subtypes involved in the interaction, and the interaction scores for each patient in full viable single-cell data. (**E**) Violin plots showing the proportions of myeloid-T cell doublets in CD45^+^-cell sorted data. Statistical significance was assessed using Student’s t-test comparing Rs versus NRs. (**F**) Selected signature scores computed in doublets involving CD8^+^ T cells and myeloid cells (left), or CD8^+^ T cells and B cells (right). Statistical significance was assessed using Student’s t-test comparing Rs versus NRs. (**G**) Examples of specific cell subsets involved in the doublet deconvolution according to clinical response. Statistical significance was assessed using Student’s t-test comparing Rs versus NRs. (**H)** Cell mutual distance analysis: Scheme and formula depicting how the distance between two different cell types was computed. (**I**) Example images of mIF staining for the indicated markers in tumors. (**J)** Heatmap showing the differences in the frequency of mutual neighbors between cell types at a 20μm neighboring radii. The heatmap display directed *p*-values (in -log_10_ and using Student’s t-test) for pairwise comparisons with red colors indicating higher frequencies in Rs and blue colors indicating higher frequencies in NRs. (**K**) Representative example images showing the density of the frequency of mutual neighbors between CD8^+^/PD-1^+^ cells and CD11c^+^ cells according to clinical response. (**L)** Violin plots displaying frequency of mutual neighbors between the indicated cell types, according to clinical response and split by stromal or intratumoral areas. Statistical significance was assessed using Student’s t-test comparing Rs versus NRs.

To increase resolution in the interactome, we focused our analysis on five pathways that emerged in the analyses of the T, B and/or myeloid compartment: complement, interferon, chemokines, interleukins and costimulation/coinhibition (**Table S6**). Again, we inferred a significantly higher number of putative interactions between TILs and myeloid cells, TILs and tumor cells, and between CD4^+^ and CD8^+^ TILs in Rs (**Fig. 5B**, **Fig. S5B)**. We inferred multiple interactions between TIL subsets predicted to be tumor-specific – including exhausted, ISG and EM-like cells – with several macrophage subsets and DCs (**Fig. 5C**). Terminally exhausted and ISG cells were predicted interacting predominantly with *C1Q*^+^ and *CXCL9*^+^ macrophages, while Pex and EM-like cells also matched against *TREM2*^+^ and *S100A8*^+^ macrophages. Based on available molecules, *CXCL9*^+^ macrophages could attract exhausted CD8^+^ TILs expressing *CCR5*, *CXCR3* and *CXCR5* through *CXCL9/10/11*, *CCL5* and *CXCL13*, respectively, and could establish adhesive interactions with them through *ICAM1* and *ICAM2*, expressed on macrophages, and *ITGB2* and *ITGAL*, on TILs (**Fig. 5D**). Through such interactions *CXCL9*^+^ macrophages could establish important niches, as seen in ovarian cancer (*29*), to support Tex function through CD28 and other costimulatory ligands. Further chemotactic and stimulatory interactions could be mediated locally by complement components, since *C3* was upregulated in exhausted TILs while *C3AR1* was expressed by complement-positive and *CXCL9*^+^ macrophages (**Fig. 5D**). Indeed, T cells upregulate and release C3 upon TCR engagement and CD28 costimulation, which can drive APC maturation and further T-cell costimulation (*45, 46*). Interestingly, *C1Q*^+^ macrophages expressed IL-15, which would provide key survival and stemness-promoting signals to local TILs in such putative niches through *IL15RA-IL2RG* (*47*) (**Fig. 5D**). Strikingly, we could not predict any crosstalks between immune cells in NRs, with the only dialogue predicted being between tumor cells and *S1008A*^+^ or *TREM2*^+^ macrophages, or DC2, mostly through complement components (**Fig. 5B-D**, **Tables S5,7**).

We sought to validate these assumptions by specifically examining physical cell doublets identified during single-cell analyses. We detected doublets occurring between T and myeloid cells (**Fig. 4A**), B and T cells (**Fig. S4A**) as well as CD4^+^ and CD8^+^ cells (**Fig. S5C**). In line with the above inferred crosstalks (**Fig. 5B-D**), we found a significantly higher frequency of T:myeloid cell doublets and a similar trend in the frequency of T:B and CD4^+^:CD8+ T-cell doublets in Rs (**Fig. 5E**, **Fig. S5C**). Importantly, in myeloid:T-cell doublets from Rs we detected overexpression of T-cell exhaustion along with CD28 costimulation signatures, implying that exhausted CD8^+^ TILs engaged in these doublets receive CD28 costimulation from the cognate myeloid cells (**Fig. 5F**). Similarly, B:T-cell doublets exhibited higher expression of T-cell costimulatory and tumor-reactivity signatures, implying that B cells could also provide CD28 costimulation to pairing tumor-specific CD8^+^ TILs (**Fig. S5D**). Finally, CD4^+^:CD8^+^ T-cell doublets overexpressed scores of cytotoxicity and tumor reactivity implying important crosstalks between tumor-specfic CD8^+^ TILs and cognate CD4^+^ helper cells (**Fig. S5D**). Similar interactions have been implicated in preventing CD8^+^ T cell exhaustion during chronic mouse infections (*48*), and could be also highly relevant for ACT. Importantly, by deconvoluting cell states involved in cell doublets we found that T:myeloid doublets were indeed enriched specifically in *CXCL9*^+^ macrophages and CD8^+^ Pex or Tex cells (**Fig. 5G**, **Fig. S5E**). Thus, these key cell crosstalks are prevalent between tumor-reactive CD8^+^ TILs and properly polarized macrophages, and are likely critical to the recruitment and functional support of these tumor-specific CD8^+^ T cells, explaining their strong association with response to TIL-ACT. Conversely, T:myeloid doublets were rare in NRs and were mostly enriched in NK-like CD8^+^ T cells and DC2 (**Fig. 5G**, **Fig. S5E**).

Surmising that cell doublets represented true physical pairs due to adhesive interactions that persisted upon tissue dissociation, we looked for these cell associations in baseline tumors. We used mIF to identify CD8^+^PD-1^+^ or CD8^-^PD-1^+^ TILs, CD11c^+^ or CD68^+^ myeloid cells and CD19^+^ B cells, and examined their total cell frequencies (**Fig. S5F)** as well as their mutual distances *in situ* (**Fig. 5H-J**). We detected higher levels of cell clusters in Rs compared to NRs, in particular revealing neighborhoods between CD8^+^ or CD8^+^PD-1^+^ TILs and CD11c^+^ cells, found both in tumor islets and in the stroma compartment (**Fig. 5J-L**, **Fig. S5G-H).** Furthermore CD8^+^PD-1^+^:CD68^+^, CD8^+^PD-1^+^:CD19^+^ or CD8^+^PD-1^+^:CD8^-^PD-1^+^ (i.e. CD4+ T cells) pairs were also higher in the stroma of Rs (**Fig. 5J**, **Fig. S5G**). These analyses collectively revealed dense immune cell networks in the TME of Rs at baseline, unlike in NRs. These intratumoral T cell:myeloid niches and other T-cell networks likely recruit and nurture tumor-reactive TILs and are at the basis of generating effective TIL therapy.

### Effective ACT-TIL therapy reprograms myeloid populations and reconstitutes antitumor CD8^+^ TIL-myeloid cell networks

Next, we sought to determine how TIL-ACT affects TME dynamics. We used bulk RNAseq data from tumors at baseline (T0) and biopsies acquired 30 days post ACT (T30) (**Fig. S1C**) to capture changes in the TME of Rs vs. NRs. Whereas ERBB2, ERBB4 and PI3K signaling pathways were downregulated in Rs, they were upregulated in NRs possibly reflecting tumor cell expansion or adaptation (**Fig. 6A**, **Fig. S6A**, **Table S8**). Importantly, responding tumors exhibited an increase in numerous inflammatory signatures (alternative complement activation, scavenger receptors, type-I interferon, TLR3, NF-κB, IL-18 and IL-10 signaling), T-cell activation (CTLA-4, PD-1, TCR, CD28, IL-2, and FasL signaling) and B-cell activation (BCR signaling), denoting reinvigoration of both adaptive and innate immunity in the TME 30 days post successful TIL-ACT. Notably, these pathways were already lower at baseline in tumors of NRs relative to Rs (see also **Fig. S3K**), and were lost in NRs’ T30 biopsies (**Fig. 6A**).

**Figure 6:**
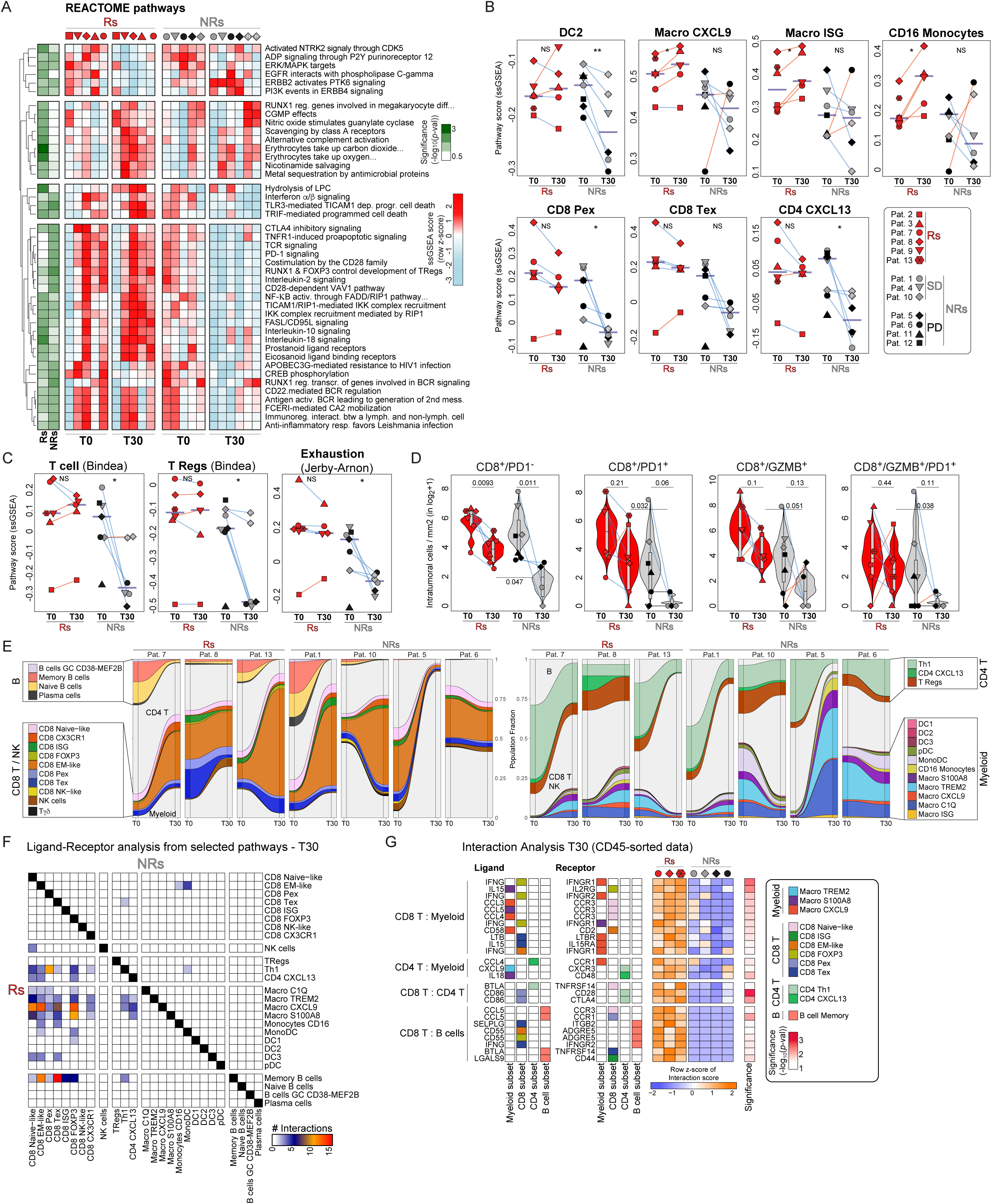
Dynamics of TME and TIL-TME interactions associated to clinical response before and after TIL-ACT. (**A**) Heatmap showing pathways that progress differently from T0 to T30 between Rs and NRs. The heatmap displays ssGSEA scores for Reactome pathways that were found either significant between T0 and T30 in Rs or NRs. Statistical significance was assessed using linear modeling as described in the methods. (**B**) Violin plots showing single-cell CD45^+^-cell sorted population signatures scores (computed by ssGSEA) in the bulk RNA sequencing data split by clinical response and time when the biopsy was taken (T0 and T30). The signatures used correspond to the 30 most significantly upregulated genes in each cell population (supplementary Table 4). Statistical significance was assessed using Student’s t-test comparing Rs versus NRs. (**C**) Violin plots showing signature scores (taken from the indicated reference) in the bulk RNA sequencing data split by clinical response and time when the biopsy was taken (T0 and T30). Statistical significance was assessed using Student’s t-test comparing Rs versus NRs. (**D**) Violin plots of cell density (in log2 (cell/mm^2^+1) between Rs and NRs at baseline (T0) and 30 days post-ACT (T30) for the indicated staining. Statistical significance was assessed using Student’s t-test comparing Rs versus NRs. (**E**) Alluvial plots displaying the progression of cell proportion of CD45^+^ cells from the baseline tumor (T0) to the tumor 30 days post infusion (T30) in CD45^+^-cell sorted single-cell data. B and CD8^+^ T cell subsets are highlighted on the left, and myeloid and CD4^+^ T cells on the right. (**F**) Heatmap displaying the number of significant ligand-receptor (from the 5 selected pathways as in Figure 5C) interactions between cell subsets in responders (lower-left part of the heatmap) and non-responders (higher-right part of the heatmap) in 30 days post-ACT biopsies (T30). (**G**) Heatmap displaying selected significant pairs of ligand-receptor, the cell subtypes involved in the interaction and the interaction scores for each patient in the CD45^+^-cell sorted data of the biopsies 30 days post-ACT.

To infer evolution dynamics in specific immune cell subets, we interrogated bulk RNAseq data at baseline and T30 employing gene signatures that discriminate for cell subtypes, which we derived from our scRNAseq data. Importantly, we observed a reconstitution post-ACT of key TME hallmarks observed at baseline, including reestablishment of CD8^+^ Pex and Tex and *CXCL13*^+^ CD4^+^ TIL signatures, and increased signatures for *CXCL9*^+^ and ISG macrophages and CD16^+^ monocytes, but uniquely in Rs. Conversely, NRs lost signatures of CD8^+^ Pex and Tex, *CXCL13*^+^ CD4^+^ TILs, Tregs and DC2 (**Fig. 6B**, **Fig. S6B)**. We validated these findings using independent immune cell gene signatures in bulk RNAseq (*49, 50*) (**Fig. 6C**), and further in paired analyses of scRNAseq at baseline and T30 from 7 patients (**Fig. S6B**). Moreover, by mIF we confirmed that responders maintained a relatively elevated frequency of CD8^+^ TILs expressing PD-1 and/or granzyme-B post-ACT, while NRs displayed significantly lower frequency of such tumor-reactive CD8^+^ TILs at T30 relative to their baseline (**Fig. 6D**). Interestingly, we observed a decrease in the frequency of intratumoral CD8^+^PD-1^-^ TILs in both Rs and NRs, representing likely bystander T cells not engaging tumor and lost due to host lymphodepletion.

We next asked whether cellular crosstalks could also be reconstituted in the TME following TIL-ACT. Analysis of the available paired scRNAseq data from baseline and T30 in seven patients (**Fig. 6E**) revealed that many of the original crosstalks were potentially restablished, with a marked increase in putative interactions predicted uniquely between the CD8^+^ and the myeloid compartment at day 30 compared to baseline, specifically in Rs (**Fig. S6C**). In these three patients, we identified again the main CD8^+^ TIL states related to tumor attack, as recognized at the steady state, i.e Pex, Tex as well as EM-like and ISG cells. Most of them were predicted to interact densely with the key myeloid subsets, especially *CXCL9*^+^ macrophages (**Fig. 6F**, **Fig. S6D**). Although most of the tumor-resident B cells were depleted post-ACT, EM-like, Tex and ISG CD8^+^ TILs were also predicted to interact with the available memory B cells (**Fig. 6F**, **Fig. S6C-D**). *IFNG* and *IFNGR1/2* interactions between CD8^+^ TIL and macrophages, respectively, implied a direct macrophage polarization by TILs leading to upregulation of *CXCL9* and *CXCL10*. The production of additional chemokines by these activated macrophages, such as *CCL3*, *CCL4* and *CCL5* explains partly how rich chemokine circuitries could help reestablish the diverse inflammatory infiltrate in these tumors post-ACT (**Fig. 6G**, **Table S9**). Conversely, as in baseline tumors, we could not predict any meaningful immune cell interactions in NRs, which were mainly predicted between CD16^+^ monocytes or monoDCs with effector-memory-like CD8^+^ TILs (**Fig. 6F-G, Fig. S6C-D**). These results suggest that successful ACT not only restored the TIL repertoire, but also reprogrammed the tumor macrophage population towards *CXCL9*^+^ cells, thereby strengthening the potential CD8^+^ T-cell:myeloid interactome.

Thus, successful TIL-ACT leads to the elimination of melanoma through the establishment of a favorable TME, with re-engraftment of tumor-reactive TILs following transfer, and improvement of the myeloid compartment after immune reconstitution. The close association of exhausted CD8^+^ TILs with CD11c^+^ myeloid cells represents a potential powerful biomarker for patient selection.

## DISCUSSION

Collectively, this study represents the most comprehensive single-cell profiling of longitudinal melanoma samples during TIL-ACT, providing insights into the tumor microenvironment and the cellular networks that are associated with clinical response to TIL-ACT.

Our comparative analysis reveals that benefit from TIL-ACT is associated with pre-existing immunity already established in the TME at the time of tumor harvest. Melanomas responding to TIL-ACT were enriched in pre-existing antigen-experienced polyfunctional intratumoral CD8^+^ T cells exhibiting features of TCR stimulation, cytotoxicity, exhaustion and CD28-costimulation (*10, 11, 29, 51*). These phenotypic traits were mostly carried by CD8^+^ Tex and Pex TILs, although the same TIL clonotypes were also distributed within the so-called ISG CD8^+^ TIL subset as well as the EM-like subset, revealing a dynamic evolution of tumor antigen-specific CD8^+^ cell states in the TME. The recently identified ISG CD8^+^ TIL subset (*25*), characterized by a dominant signature of IFN response, remains intriguing. Here, these cells emerged as antigen-experienced, with signatures of TCR activation and mild exhaustion indicating that they are likely tumor-reactive CD8^+^ cells embedded in IFN-rich tumor niches.

Our data also suggest that tumor-reactive CD8^+^ TILs are engaged in complex and meaningful cell networks in the TME at baseline, specifically in patients who subsequently respond to TIL-ACT. Myleoid cells were among the key populations contributing to these interactions, and beared unique gene signatures in responders, specifically overexpressing antigen-processing and presentation, costimulatory and complement genes as well as type-I IFN signatures and CXCL9/10 chemokines. Recent studies have demonstrated the relevance of tumor-intrinsic (*17*) and myeloid-mediated DNA-sensing STING activation for tumor IFN signaling (*52*), the recruitment of TILs and the orchestration of anti-tumor immune attack (*53*). These are hinged on the cooperation of CCL5 and CXCL9/10 chemokines, which ensure recruitment and retention of TILs in the TME by intratumoral myeloid cells (*36*) and promote response to ICB (*54*). Furthemore CXCL9/10-expressing myeloid cells are associated with the presence of CXCL13-expressing TILs (*55*). Importantly, these chemotactic interactions, along with the appropriate adhesion partners reported here, are at the basis for the formation of cell pairs between effector TILs and myeloid cell partners. In baseline melanoma responsive to ACT therapy, we predicted the cellular interactome by ligandome analysis and documented a strong intratumoral crosstalk in the chemokine, complement and interferon pathways and adhesions between tumor-reactive CD8^+^ TIL and activated myeloid cells. Although traditionally considered undesirable, doublets can provide valuable information on immune cells interactions (*56*). We validated these interactions by demonstrating physical TIL-myeloid cell doublets, and by quantifying -through mIF and niche analysis-cellular neighborhhods *in situ* in the tumor of origin, which revealed denser clusters of intratumoral T cell:myeloid niches in responders at baseline. Importantly, based on scRNAseq findings, the T cell:myeloid doublets were largely populated by Tex CD8 and IFN-activated myeloid cells, confirming the specific implication of these cell subsets in driving T-cell attack at baseline. In addition of recruiting TILs, IFN-activated myeloid cells could also endow them with costimulatory signals (*29*), thereby providing a rich repertoire of relevant polyfunctional TILs at baseline that could be subsequently expanded *ex vivo* for TIL-ACT. Importantly the TME of NRs displayed mininal numbers of those interactions and almost no cellular crosstalk when focusing in pathways such as costimulation, complement, type-I IFN and chemokines.

Interestingly, many of the above cellular crosstalks may be also implicated in driving response to ICB. Yet, four of the six responding patients in this study had failed prior ICB, revealing that additional inhibitory mediators were at play in the TME at baseline, preventing response to ICB, but could have been eliminated upon TIL-ACT. In fact, TIL reinvigoration by *ex vivo* expansion and host lymphodepletion could contribute to overcome TME suppression. Indeed, by comparing the evolution dynamics of TIL and TME states from baseline to day-30 post ACT, we observed an optimal evolution of the TME specifically in responders, where TILs with tumor-reactive features were the preponderant TIL subset and the frequency of IFN-activated myeloid cells increased, while the T-cell interactome with myeloid cells further increased and diversified, indicating that effective ACT-TIL therapy reprograms myeloid populations and increases antitumor CD8^+^ TILs-myeloid cell networks. Such interactive immune TME could sustain the persistence and functionality of *ex vivo* reinvigorated and adoptively transferred T cells via additional costimulatory signals. In addition, homotypic CD4:CD8 doublets and B:TIL doublets, identified uniquely in responders, could provide complementary costimulatory supporting signals.

Our detailed TME states and dynamics as well as cell-cell interactome description will facilitate the development of biomarkers of response and guide the next generation of adoptive cell-based immunotherapies to achieve maximal clinical benefit. Our findings suggest that tissue signatures documenting the presence of intratumoral tumor-reactive TIL:myeloid niches with traits of polyfunctionality, fitness and costimulation can be utilized to select for patients that can maximally benefit from TIL-ACT approaches with traditional IL-2-based TIL expansion methodologies.

## Acknowledgments

We are grateful to the patients and their families for their dedicated collaboration.

We thank Virginie Zimmer, Jade Poll, Jean-Paul Rivals and Philippe Gannon from the Center of Experimental Therapeutics (CTE) Biobank for their assistance.

We thank Christophe Sauvage for assistance with single cell encapsulation and cDNA library preparations; the Agora Flow Cytometry Facility of the University of Lausanne and the Lausanne Genomic Technologies Facility for scRNAseq/scTCRseq and RNA-seq analysis.

We also apologize to colleagues whose work could not be cited because of space limitation.

## Funding

In the last three years G. Coukos has received grants, research support or has been coinvestigator in clinical trials by Bristol-Myers Squibb, Tigen Pharma, Iovance, F. Hoffmann-La Roche AG, Boehringer Ingelheim. The Lausanne University Hospital (CHUV) has received honoraria for advisory services G. Coukos has provided to Genentech, AstraZeneca AG, EVIR. Patents related to the NeoTIL technology from the Coukos laboratory have been licensed by the Ludwig Institute, on behalf also of the University of Lausanne and the CHUV, to Tigen Pharma. G. Coukos has previously received royalties from the University of Pennsylvania for CAR-T cell therapy licensed to Novartis and Tmunity Therapeutics.

Dr. Dangaj Laniti reports grants from Swiss National Foundation (SNF) R’Equip (316030_205644).

## Author contributions

Conceptualization: DB, EG, AO, SZ, DDL, GC.

Methodology: DB, NF, JC, SB, DL, UD, AG, MM, AB, ST, ZT, CA, IC, JCO, FB, RG, MP, DDL, GC.

Investigation: DB, EG, NF, JC, JD, FB, IC, MBS, AB, ST, DDL, GC.

Visualization: DB, EG, JD, FB, ST, DDL

TIL production process: PG, LK

Sample Collection and Coordination: RaD, VG, JP, JPR, LK

Funding acquisition: DDL, GC

Patients treatment: EG, AO, FH, RaD, ReD, MI, MOdO, BN, KH, SZ, MM, VA, CD, NS, JOP, ND, OM, LT, GB, RD

Supervision: DDL, GC, CS, AH, LT

Writing – original draft: DB, EG, DDL

Writing – review & editing: DB, EG, DDL, GC

### Competing interests

SZ is currently an employee of F. Hoffmann-La Roche.

RD has intermittent, project focused consulting and/or advisory relationships with Novartis, Merck Sharp & Dhome (MSD), Bristol-Myers Squibb (BMS), Roche, Amgen, Takeda, Pierre Fabre, Sun Pharma, Sanofi, Catalym, Second Genome, Regeneron, Alligator, T3 Pharma, MaxiVAX SA, Pfizer and touchIME outside the submitted work.

OM has consulting/advisory roles for Bristol Myers Squibb, MSD, Roche, Novartis, Amgen, Pierre Fabre, Neracare; research grants from Bristol Myers Squibb, MSD, Amgen. PCL; consultant advisor or paid speaker for Bristol Myers Squibb, MSD, Novartis, Pierre Fabre, Amgen, Nektar; has received research funding from Bristol Myers Squibb, Pierre Fabre.

GC has received grants from Celgene, Boehringer-Ingelheim, Roche, Bristol Myers Squibb, Iovance Therapeutics and Kite Pharma. The institution G.C. is affiliated with has received fees for G.C.’s participation on advisory boards or for presentation at a company-sponsored symposium from Genentech, Roche, Bristol Myers Squibb, AstraZeneca, NextCure, Geneos Tx and Sanofi/Avensis. G.C. holds patents around antibodies and receives royalties from the University of Pennsylvania regarding technology licensed to Novartis.

DDL, AH, and GC are inventors on patent applications filed by the Ludwig Institute for Cancer Research Ltd pertaining to the subject matter disclosed herein and such patent applications have been licensed to Tigen Pharma SA.

All the other authors have no conflict of interest to declare.

### Data and materials availability

Single-cell and bulk RNA sequencing data will be made publicly available in the Gene Expression Omnibus (GEO) at the time of publication.

## Materials and Methods

### Patients

From March 2018 to January 2021, we enrolled fifteen patients in a single-center phase 1 investigator-initiated trial designed to test the feasibility of ACT with TILs (ClinicalTrials.gov NCT03475134). Among them, two patients did not receive any trial treatment: one patient failed during the rapid expansion phase (REP)-TIL and another one withdrew informed consent from the study before the start of the treatment. Thirteen eligible patients constitute the ‘per protocol’ cohort of the study for efficacy analysis (**Table S1**). Eligible patients were adults with histologically proven unresectable locally advanced (stage IIIc) or metastatic (stage IV) melanoma who have progressed on at least 1 standard first line therapy, including but not limited to chemotherapy, BRAF and MEK inhibitors, anti-CTLA4, anti-PD-1 or anti-LAG3 antibodies and/or the combination. When feasible, a biopsy of one metastatic deposit was performed at screening to assess and quantify the intratumoral CD3^+^ and CD8^+^ infiltration by a dedicated Pathologist (JD). In some cases, archived FFPE material from diagnosis was retrieved for analysis. Patients were required to have an accessible metastasis to procure for TILs with acceptable anticipated perioperative risk and also at least one separate additional measurable tumor lesion on CT. Patients were required to have a good general health status (ECOG PS ≤ 2), sufficient cardiopulmonary function, including a cardiac stress test showing no reversible ischemia; adequate respiratory function with forced expiratory volume in 1 second (FEV1) ≥ 65% predicted, forced vital capacity (FVC) ≥ than 65% predicted and DLCO ≥ than 50% predicted corrected; and left ventricular ejection fraction (LVEF) > 45%. Patients with active brain metastases, autoimmune conditions or acquired immunodeficiency were excluded. Patients were required to have adequate normal organ and marrow function, defined as hemoglobin ≥ 8 g/100 ml; absolute neutrophil count ≥ 1.0 × 10^9^ (≥1,000 mm^−3^); platelet count ≥100 × 10^9^ (>100,000 mm^−3^); serum creatinine ≤ 1.5× of the institutional upper limit of normal; and AST and ALT ≤ 3× of the institutional upper limit of normal. Patients with symptomatic and/or untreated brain metastases were excluded. Patients with definitively-treated brain metastases will be considered for enrollment, as long as lesions are stable for ≥ 14 days prior to beginning the chemotherapy, there are no new brain lesions, and the patient does not require ongoing corticosteroid treatment. Bridging-therapy was allowed as per investigator-choice and upon discussion with the Principal Investigator.

Clinically eligible patients underwent surgery for TILs harvest and *ex vivo* expansion. Only patients having sufficient numbers of pre-REP TILs (TIL numbers > 50 millions) were offered to receive TIL-ACT treatment. TILs were successfully expanded for all 13 patients from tumor deposits resected by surgery and the median number of TILs infused was 55.0 billions of cells (range: 12.8-84.7).

### Study design

The primary endpoints were feasibility and safety of ACT using autologous TILs in our center. Key secondary endpoints were feasibility and safety of nivolumab rescue following TIL-ACT, clinical efficacy of the treatment with respect to ORR, PFS, according to RECIST v1.1 and OS. Overall Survival (OS) was defined as the time from start of NMA chemotherapy until death from any cause, for a maximum of 5 years. If there is no death date, the patient is censored on the last day known to be alive. Exploratory objectives included collection of exploratory translational data regarding the biological effects of the TIL-ACT and its interaction with the tumor microenvironment, using paired tumor biopsies before and after treatment, as well as blood samples. Tumor samples were collected at screening (if feasible), at surgery (tumor material for pre-REP), at day 30 after TILs-ACT, after 4 weeks of nivolumab treatment if applicable (optional) and at progression (optional). Detailed list of patients’ samples available for translational analysis is provided in **Figure S1C**. Adverse events were recorded according to NCI Common Terminology Criteria for Adverse Events (CTCAE v5.0).

### Study conduct

Patients received lymphodepletion regimen consisting of fludarabine (25 mg/m^2^/day) for 5 days and cyclophosphamide (60 mg/kg/day) for 2 (overlapping) days, followed by the infusion of T lymphocytes, which is followed by the administration of intravenous boluses of high dose IL-2 (720,000 IU/kg) starting 3 hours post-TIL infusion, then every 8h at minimum counting from the start of each administration, for a total of 8 doses maximum, with a maximum interval of 24h. In particular, ten patients (76.9%) received a full course of lymphodepleting chemotherapy (cyclophosphamide and fludarabine) without dose modification. All patients initiated high-dose IL-2 treatment and received a median of 5 doses of IL-2 (range 1-8) (**Table S1**). Adverse effects were primarily attributable to lymphodepletion and IL-2 administration. Common non-hematologic adverse events included nausea, hypophosphatemia and capillary leak syndrome (hypoalbuminemia, weight gain and pulmonary edema).

Following day 30, at the completion of the TLT period, eligible patients will be started on nivolumab, as long as no TIL-ACT related toxicity > to grade 1 is observed. Nivolumab, at a dose of 240 mg IV every two weeks will be administered for the first 12 months, followed by nivolumab at a dose of 480 mg IV every four weeks for the next 12 months until unacceptable toxicities or confirmed disease progression.

Only 3 out of 11 patients (27%) were eligible for nivolumab rescue. Patient #7 progressed at 6 months after TILs infusion (Nov 2019) with new inguinal lymph nodal lesions and received six cycles of nivolumab treatment with initial stability of the disease and then new PD for which TKI therapy was started (Sept 2020) and still ongoing. Patient #9 progressed five months after TILs infusion and received five cycles of nivolumab with a further progression after three months. Patient #12 progressed at first-month assessment and received six cycles of nivolumab with further PD.

### Synthetic Control Arm (SCA)

The SCA for our analysis is based on historical, real-world data from CHUV (Centre hospitalier universitaire vaudois) patients, taking into account the current treatment strategies and the consistency of characteristics with the ATATIL patients. In order to build a synthetic comparator arm for patients who failed standard of care we performed a search in our institutional clinical research data warehouse (June 2021) complemented by a number of fields curated specifically for our melanoma cohort. Uveal melanoma and patients who took part of an ACT-TIL trial were excluded. Patients who have progressed on first line PD-1 or PD-1 + CTLA-4 blockades and, for BRAF V600 mutated melanoma, who subsequently failed BRAF + MEK inhibitors were identified via a structured search by a physician. The BRAF status of the tumors has been used as a stratification factor. Of note, in the present analysis BRAF-positive subgroup refers precisely to BRAF-V600 mutated melanoma (potentially treated by BRAF inhibitors), while the BRAF-negative subgroup includes “non-mutated BRAF-V600” patients.

### Immunohistochemistry, TIL assessment and scoring

For each case, representative tumor slides with the most viable component and higher inflammatory infiltrate were selected for immunohistochemical (IHC) staining on formalin-fixed paraffin-embedded (FFPE) whole tissue sections. CD3 (2GV6, CONFIRM, Rabbit Monoclonal, Roche, Basel, Switzerland), CD8 (C8/144B, Mouse Monoclonal, Dako, Glostrup, Denmark), PD-1 (NAT105, Mouse Monoclonal, CellMarque, Rocklin, United States) and CXCL13 (BLC/BCA-1, Goat Polyclonal, IgGR&D Systems, Minneapolis, United States) expressions were assessed. Briefly, immunohistochemical staining (IHC) was performed using the Ventana Benchmark Ultra (Roche Ventana Tucson, Arizona, USA) for the CD3, CD8, PD-1 and CXCL13 stains, by following the manufacturer’s instructions. Four µm-thick FFPE sections were subjected to routine deparaffinization, rehydration, and antigen retrieval procedures. Each section was incubated with the primary antibody, followed by the secondary antibody. The ultraView Universal DAB Detection Kit for CD3, CD8, and CXCL13 IHC (Ref: 05269806001, Roche Ventana); and the OptiView DAB Detection Kit for the PD-1 IHC (Ref: 06396500001, Roche Ventana) were used as detection system. Tissue counterstaining was performed with Hematoxylin from Gil II solution (Ref: 105175, MERCK). Sections of human tonsil were used as positive control. Evaluation was performed independently by one pathologist (JD) without knowledge of clinical information.

For each sample, at least 10 high-power fields (HPF) with a diameter of 500 µm were selected. These fields were distributed over the whole tumor sample. Intra-tumoral T lymphocytes were qualitatively assessed by CD3 (low to high), along with spatial distribution (stroma *vs.* tumor) and heterogeneity. CD8/CD3 ratio and CD8 ranges per HPF were then evaluated. The final TILs score was the mean intratumoral CD8^+^ cells in at least 10 HPFs. The tumor-infiltrating CD8^+^ T-cells were evaluated and classified as “intratumoral” if they were in direct contact with tumor cells. Cells stained positive in the stromal compartment and within the borders of the invasive tumor or in areas of necrosis were not evaluated. For cases with significant region variations in lymphocyte distribution, each region was evaluated separately and an average value was assessed in > 10 HPF whenever possible. The number of PD-1 (membranous immune cell labeling) and CXCL13 (cytoplasmic and perinuclear cell labeling) positive cells in the same representative high-power fields (HPF) were also assessed and averaged.

### Multispectral immunofluorescence tissue staining and image analyses

For the multiplexed staining, FFPE sections were stained by an automated immunostainer (DISCOVERY ULTRA, Ventana Roche). First, the heat-induced antigen retrieval in EDTA buffer (pH 8.0) was performed for 92 min at 95°C. Multiplex staining was performed in consecutive rounds, each round consisting of protein blocking, primary antibody incubation, secondary HRP-labeled antibody incubation, OPAL detection reagents and then antibodies heat denaturation. The Multiplex IF images were acquired on Polaris imaging system (Perkin Elmer). Tissue and panel specific spectral library of the specific panel individual fluorophore and tumor tissue autofluorescence were acquired for an optimal IF signal unmixing (individual spectral peaks) and multiplex analysis. The IF stained slides were pre-scanned at 10x magnification using the Phenochart whole-slide viewer. Using the Phenochart™ whole-slide viewer, regions of interest (ROI) representative of the all sample were adquired. InForm 2.5.1 software was used for training and phenotyping analysis. The images were first segmented into specific tissue categories of tumor, stroma and no tissue, based on the cytokeratin and DAPI staining using the inForm Tissue Finder™ algorithms. Individual cells were then segmented using the counterstained-based adaptive cell segmentation algorithm. Quantification of the immune cells was then performed using inForm active learning phenotyping algorithm by assigning the different cell phenotypes across several images chosen for the project. IF-stained cohorts were then batch processed on the whole image and data were exported via an in-house developed R-script algorithm to retrieve every cells’ x,y coordinates and staining positivity and intensity. To calculate cell densities of different populations we counted the number of specific cells phenotypes in both tumor and stroma compartment on a whole-slide basis. Counts of each cell-type (tissue-specific) were divided by the area of tissue (mm^2^) to obtain a density (number of cells/mm^2^).

### Neighborhood cell-to-cell distance analysis and density map

Given two sets of points in a two dimensional space we could implement a measure that estimate their vicinity. Given the set A={A_1_, …., A_N_}c, and the set B={B_1_, …, B_K}_, we express the coordinate of each point A as x_an_, y_an_, and for each point B, x_bn_, y_bn_.

For each A we measure the number of neighbors of type B in a distance D < ε, where D is:

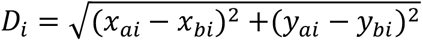

This can be graphically view as counting the number of neighbors in a circle centered on the point Ai, with radius ε. Therefore, we can define the function “S” that, applied to an Ai, gives the number of neighbors of type B around this element.

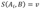

The extremes of the function S goes from v = 0 to “K”, where 0 means no neighbors for an element of A, and “K” means that all the elements of the set B can be found below the distance ε from that specific Ai.

From the function “S”, we can create another function:

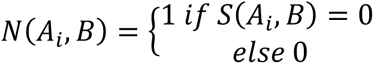

We can apply the previous equation in a summation to get number of element A that have at least one neighbor.

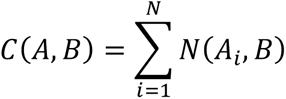

Once defined the function for the set “A”, this function can be reversed to count the number of neighbors of type A around a given element of the set B.

Indeed, to take into account the interaction between A to B, and the interaction between B to A, we can define another function “M” (that we can call mutual or mutual frequency):

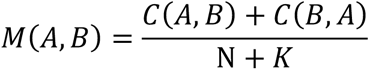

We can then extend this function to multiple sets and apply these metrics to any kind of pair, changing starting point/ending point and the radius-diameter of interest (20μm, 45μm, 100μm). Using precise cells coordinates and phenotypes it is possible to digitally represent them across the tissue. Particularly, to visualize densities of niches in the digital reconstructed tissue, we can use the coordinates of such niches and a 2D Kernel Density Estimation (KDE). The KDE allows us to approximate the density distribution of specific type of niches (or any other kind of species in a 2D surface) and shows the equiprobability lines for such densities. It is important to notice that, the highest is the number of niches, the highest will be the overlap between the equiprobability lines and the actual tissue.

### Tissue processing

Resected tumors were chopped into 1-2 mm^2^ pieces and, along with post-infusion biopsies, cryopreserved in 90% human serum + 10% dimethyl sulfoxide (DMSO) and additional pieces were snap frozen for bulk RNA extraction. For single-cell experiments, both frozen and fresh material were used as starting material. PBMCs were isolated from blood collected in EDTA tubes and cryopreserved in 90% human serum + 10% DMSO.

The day of the assay, pathofrozen pieces were thawed in RPMI + 10% FBS and chopped in small pieces using a scalpel. Tissue was dissociated in RPMI + 2% Gelatin (#G7041, Sigma-Aldrich) + 200 IU/mL Collagenase I (#17100-017, ThermoFisher Scientific) + 400 IU/mL Collagenase IV (#17104-019, ThermoFisher Scientific) + 5 IU/mL Deoxyribonuclease I (#D4527, Sigma-Aldrich) + 0.1% RNasin Plus RNase Inhibitor (#N2618, Promega) for 15-30min (depending of sample size and consistency) at 37°C and shaken at 160rpm. Digested cells were filtered using a 70 μm strainer and resuspended in PBS + 1% Gelatin + 0.1% RNasin. Cells were manually counted with hematocymeter then stained for viability with 50uM/mL of Calcein AM (#C3099, Thermo Fisher Scientific) and FcR blocked (#130-059-901, Miltenyi Biotec) for 15min at RT. After incubation and washing, cells were stained with CD45-APC (#304012, BioLegend) for 20min at 4°C. After washing, cells were resuspended in PBS + 0.04% BSA (Sigma-Aldrich) + 0.1% RNasin and DAPI staining (Invitrogen) was performed.

### Bulk RNA sequencing library preparation and processing

Total RNA was extracted from snap frozen tissues. Samples were lyzed in TRIzol reagent (Invitrogen) using a tissue lyzer (Qiagen) and the RNA was purified with the RNeasy kit (Qiagen). RNA quality was assessed with a Fragment Analyzer (Agilent) and Nanodrop spectrophotometer (Thermofisher). Quantification was performed with the Qubit HS RNA assay kit (InVitrogen). RNA sequencing libraries were prepared using the Illumina TruSeq Stranded Total RNA reagents according to the protocol supplied by the manufacturer and sequenced using HiSeq 4000.

Illumina paired-end sequencing reads were aligned to the human reference GRCh37.75 genome using *STAR* aligner (version 2.6.0c) and the 2-pass method as briefly follows: the reads were aligned in a first round using the *--runMode alignReads* parameter, then a sample-specific splice-junction index was created using the *--runMode genomeGenerate* parameter. Finally, the reads were aligned using this newly created index as a reference. The number of counts was summarized at the gene level using *htseq-count* (version 0.9.1). The Ensembl ID were converted into gene symbols using the *biomaRt* package and only protein-coding, immunoglobulin and TCR genes were conserved for the analysis. Read counts were normalized into reads per kilobase per million (RPKM) and log2 transformed after addition of a pseudo-count value of 1 using the *edgeR* R package. As the data came in three different batches, we applied a batch correction algorithm using the *ComBat* function of the *sva* R package by using the patient origin as a covariate in the model.

### FACS sorting, encapsulation and library construction

40’000 CD45^+^ cells and 40’000 total live cells were sorted on a MoFlo Astrios (Beckman Coulter) and collected in separated 0.2mL PCR tubes containing 10μl in PBS + 0.04% BSA + 0.1% RNasin. After sorting, cells were manually counted with hemocytometer and viability was assessed using Trypan blue exclusion. *Ex vivo* CD45 cells from tumor were resuspended at a density of 600-1200 cells/μl when possible with a viability of >90% and subjected to a 10x Chromium instrument for single-cell analysis. Single-cell RNA libraries were generated using the Chromium Next GEM Single Cell 5’ Library and Gel beads kit v1.1 for the CD45-sorted population or using the Chromium Next GEM Single Cell 3’ Library and Gel beads kit v3.1 for the live-sorted cells, according to the manufacturer’s instructions. For each sample, 15’000 cells were loaded into the Chromium machine when possible (i.e. the total of sorted cells if <15’000), encapsulated and barcoded following the manual (CG000388 for 3’ or CG000207 for 5’ technology). After encapsulation and reverse transcription, 14 PCR cycles were used to amplify cDNA. All library construction steps were performed according to the manufacturer’s protocol. For each sample, 3’GEX, 5’GEX and 5’VDJ enriched libraries were generated. Complementary DNA and library quality were examined on a Fragment Analyzer (Agilent) and quantification was performed with the Qubit HS dsDNA assay kit (InVitrogen).

For the FACS sorting of post-ACT biopsies, depending of the percentage of CD45^+^ fraction in the sample, two sorting strategies were adopted. If the CD45^+^ fraction was inferior to 75% of total viable cells, 40’000 CD45^+^ and 40’000 CD45^-^ viable cells were sorted in two separated tubes. If the CD45^+^ fraction was superior to 75% of total viable cells, 40’000 total viable cells were collected in one tube. All cells were sorted on a MoFlo Astrios (Beckman Coulter) and collected in separated 0.2mL PCR tubes containing 10μl in PBS + 0.04% BSA + 0.1% RNasin. After sort, cells were manually counted with hemacytometer and viability was assessed using Trypan blue exclusion. If CD45^+^ and CD45^-^ were sorted separately, cells were mixed with a ratio of 70% CD45^+^ and 30% CD45^-^. All cells were resuspended at a density of 600-1200 cells/mL if possible with a viability >90% and subjected to a 10x Chromium instrument for single-cell analysis. Only the CD45^+^ compartment was analyzed in this study.

### Alignment, filtering and processing of scRNA-seq libraries

We analyzed the following scRNAseq GEX/TCR datasets: 1) 3’GEX from baseline sorted viable cells (total TME dataset) of 10/13 patients which retained cell stoichiometry of the total TME, 2) 5’GEX from baseline CD45^+^-sorted cells of 13/13 patients which permits relative and deep phenotyping of only immune cells, including rare populations and 3) 5’GEX from CD45^+^-sorted cells of 7/13 patients 30 days post TIL-ACT which enabled us to track the dynamics of immune cell infiltration post TIL-ACT. 4) 5’ scTCR-seq (VDJ) data from baseline CD45^+^-sorted cells of 13/13 patients (same libraries than 5’ GEX).

The scRNA-Seq reads were aligned to the GRCh38 reference genome and quantified using *cellranger* count (10X Genomics, version 4.0.0). Filtered gene-barcode matrices that contained only barcodes with unique molecular identifier (UMI) counts that passed the threshold for cell detection were used and processed using the *Seurat* R package version 4.0.1. Two different Seurat objects were created: one containing CD45^+^-sorted cells from tumors (15 baseline tumor samples from 13 patients and 7 samples for post-ACT tumors from 7 patients using 5’ sequencing technology) and one containing all viable cells from baseline tumors (3’ technology of 12 samples from 10 patients using 3’ sequencing technology (see **Figure S1C** for sample availability). The baseline tumor samples involving two different sites for the same patient (patients 10 and 13) were pooled for subsequent analyses unless otherwise mentioned.

For CD45^+^-sorted cells from tumor data (5’GEX), we defined and removed low-quality cells containing more than 10% of mitochondrial reads. A table summarizing the number of cells per sample before and after filtering appears below. For CD45^+^-sorted cells from tumor data, the number of genes expressed per cell averaged 1’240 (median: 1’458) and the number of unique transcripts per cell averaged 4’206 (median: 2’926). For sorted viable cells from tumor data (3’GEX), the number of genes expressed per cell averaged 2’136 (median: 1’811) and the number of unique transcripts per cell averaged 7’297 (median: 4’961). We did not filter out cells from the baseline sorted viable cell dataset (3’) as this dataset was mainly built to derive the stoichiometry of cell types.

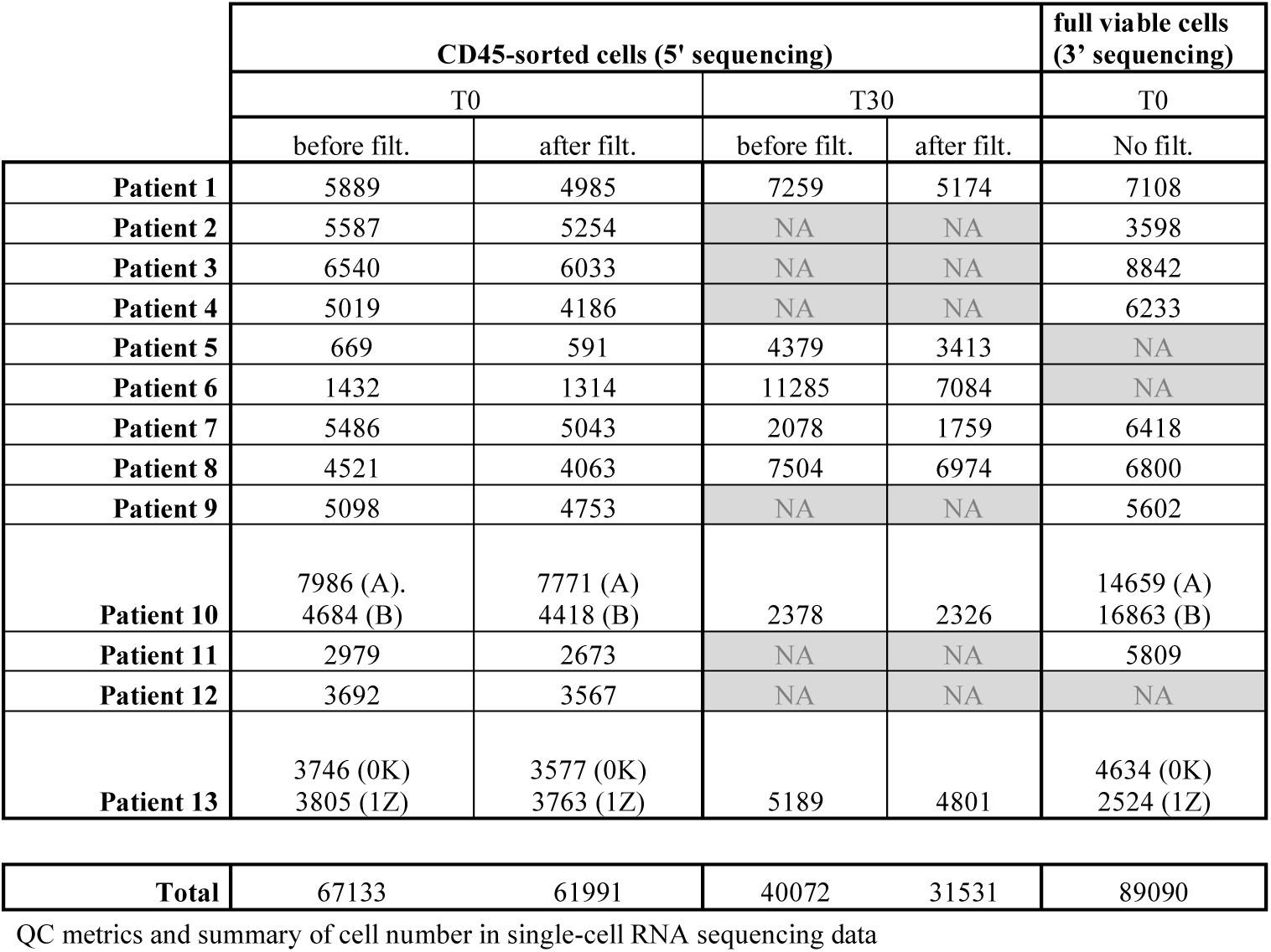

|  | CD45-sorted cells (5' sequencing) |  |  |  | full viable cells (3' sequencing) |
| --- | --- | --- | --- | --- | --- |
|  | T0 |  | T30 |  | T0 |
|  | before filt. | after filt. | before filt. | after filt. | No filt. |
| **Patient 1** | 5889 | 4985 | 7259 | 5174 | 7108 |
| **Patient 2** | 5587 | 5254 | NA | NA | 3598 |
| **Patient 3** | 6540 | 6033 | NA | NA | 8842 |
| **Patient 4** | 5019 | 4186 | NA | NA | 6233 |
| **Patient 5** | 669 | 591 | 4379 | 3413 | NA |
| **Patient 6** | 1432 | 1314 | 11285 | 7084 | NA |
| **Patient 7** | 5486 | 5043 | 2078 | 1759 | 6418 |
| **Patient 8** | 4521 | 4063 | 7504 | 6974 | 6800 |
| **Patient 9** | 5098 | 4753 | NA | NA | 5602 |
| **Patient 10** | 7986 (A).<br>4684 (B) | 7771 (A)<br>4418 (B) | 2378 | 2326 | 14659 (A)<br>16863 (B) |
| **Patient 11** | 2979 | 2673 | NA | NA | 5809 |
| **Patient 12** | 3692 | 3567 | NA | NA | NA |
| **Patient 13** | 3746 (0K)<br>3805 (1Z) | 3577 (0K)<br>3763 (1Z) | 5189 | 4801 | 4634 (0K)<br>2524 (1Z) |
| **Total** | 67133 | 61991 | 40072 | 31531 | 89090 |

QC metrics and summary of cell number in single-cell RNA sequencing data

The full baseline sorted viable cell dataset (3’) contained real stoichiometry for all cell types except for patient 1 where the scRNAseq library was enriched for malignant cells to reach an approximate rate of CD45^+^ and non-CD45^+^ cells of fifty percent each for a better characterization of the malignant compartment. Thus, for analyses involving stoichiometrical comparisons, the real stoichiometry of cell types was deduced and used by forcing the CD45^+^ cells to represents 93.9% (value obtained by FACS analysis) of the total number of cells.

scTCR-seq (VDJ) data were aligned to the same human genome using the *cellranger vdj* (10X Genomics, version 3.1.0). Only true cells (with a “True” label in the “is_cell” column of the all_contig_annotations.csv file) were kept for further analyses. Cells from the VDJ sequencing were mapped to the scRNA-Seq data (5’GEX).

### Single-cell clustering analysis

The data was processed using the *Seurat* R package (version 4.0.1) as follows: data counts were log-normalized using the *NormalizeData* function (scale.factor=10’000) and then scaled using the *ScaleData* function by regressing the mitochondrial and ribosomal rate of read contents, the number of read count per cell (nCount_RNA), and cell cycle parameters represented by S phase and G2/M phase scores (computed using the *CellCycleScoring* function with the list of genes provided internally in the *Seurat* package). Dimensionality reduction was performed using the standard Seurat workflow by principal component analysis (*RunPCA* function) followed by tSNE (*RunTSNE* function with tsne.method=”Rtsne”) and UMAP projection (*RunUMAP* function with min.dist = 0.75) using the first 75 principal components. The k-nearest neighbors of each cell were found using the *FindNeighbors* function run on the first 75 principal components (nn.eps=0.5), and followed by clustering at several resolutions ranging from 0.1 to 10 using the *FindClusters* function. Unless otherwise mentioned, default parameters were used. After the annotation of cells (see next section), the data was integrated by main cell type (CD8^+^ T cells with NK cells, myeloid cells, B cells) following the Satija lab integration procedure using anchors and as follows: expression data Seurat objects were split per sample and then normalized individually using *NormalizeData*. The most variable features were individually identified using the *FindVariableFeatures* function. The 800 best genes selected for running the integration were found using *SelectIntegrationFeatures* on all split Seurat objects together. Every Seurat objects was then scaled a PCA was run individually using only the selected features. Anchors were then found using the *FindIntegrationAnchors* function (reduction.method=”rpca” and 30 principal components) and these anchors were used for integration by using the *IntegrateData* function. This merge integrated object was then subjected to dimension reduction (20 principal components) and clustering as explained above. For B cells integration, samples with too few B cells (patient #3, #5, #6 and #11) could not be integrated and were thus removed from the data.

### Annotation of single cells

#### Main cell-type annotation

The annotation of main cell types was performed on both datasets using the combination of several different methods: i) differential gene expression of clusters at different resolutions using the Seurat *FindAllMarkers* function followed by literature curation ; ii) investigation of the expression of known canonical gene markers for melanoma cells (*MLANA*, *PRAME*, *SOX10*, *S100B*), immune cells (*PTPRC*), T cells (*CD3E*, *CD8A*, *CD8B*, *CD4*), B cells (*CD79A*, *MS4A1*), CAFs (*DCN*, *FAP*), endothelial cells (*PECAM1*, *VWF*), plasma cells (immunoglobulins), Myeloid cells (*CD68*, *HLA-DRA*, *LYZ*, *CD86*), plasmacytoid DC (*LILRA4*) ; iii) automated annotation using the *singleR* package and using gene expression centroids (average gene expression profiles per cell type) derived from several studies: Yost *et al*. (*18*), Guo *et al*. (*19*), Zhang *et al*. (*20*), Oliveira *et al*. (*9*), Zilionis *et al*. (*39*) and the human primary cell atlas (HPCA) reference as inferred in the *singleR* package. Of note, the centroids from Yost *et al*., Guo *et al*. and Zhang *et al*. were taken from Wu *et al*. (*57*).

#### Malignant cell annotation

The malignant cell annotation from the sorted viable cell dataset was also confirmed by using two other methods: (i) We derived a signature of genes specific to melanoma using pan-cancer normalized bulk RNA sequencing data from The Cancer Genome Atlas (TCGA) (https://gdc.cancer.gov/about-data/publications/pancanatlas). We performed differential gene expression using the regularized linear model as implemented in the *limma* R package to extract the 50 most discriminant genes between melanoma (SKCM) and all other cancer types together. We computed a melanoma specific score using the *AUCell* R package to ensure that private clusters found in the sorted viable cell data were melanoma using they were significantly higher than score of normal CD45^+^-positive cells. (ii) We inferred the copy-number variation (CNV) using *CopyKAT* R package. For computational optimization, we subsampled the sorted viable cell dataset by randomly selecting 200 cells from each cluster at resolution 0.1 then we run *CopyKAT* by using the normal cells (immune, stromal and endothelial) as reference. Inferred CNV profiles of 200 randomly selected cells per cluster exhibited clear CNVs, indeed confirming that these cells were of malignant origin. The copy-number alterations (CNA) were extracted from the *CopyKAT* output and represented as heatmaps by sorting genes per chromosomal location and cells per cell type using *pheatmap* R package. We then computed the number of genes falling in amplified (log ratio of the CNV > 0.2) or deleted (log ratio of the CNV < -0.2) per patient. To do so, we extracted genomic locations of CNA regions as given by *CopyKAT* output and loaded the ‘full.ann’ database consisting of genomic locations annotated with gene symbols. We then computed the overlap between amplified and deleted regions and the ‘full.ann’ database using the *findOverlaps* function from the *IRanges* R packages. The number of genes that were deleted, amplified or the sum of both (all CNV) per patient was then plotted using the *violin_plot* function from the *plotrix* R package. We also checked if deleted and amplified genes were cancer driver genes by intersecting the gene list with a list of 204 oncogenic and tumor suppressor cancer driver genes specific for melanoma or pan-cancer that were extracted from Bailey *et al*. (*58*).

#### High-resolution cell type annotation

Once major clusters were annotated for both datasets (malignant, T cells, B cells, myeloid cells, CAFs and endothelial cells), we performed detailed and curated cell annotation on the CD45^+^-sorted single-cell dataset which has higher resolution of the tumor microenvironment and then projected transcriptomics profiles on the sorted viable cell dataset for annotation as explained below.

#### High-resolution cell type annotation of T cells and NK cells

For finer annotation of the T/NK cells, the cells were first classified as CD8^+^, CD4^+^, double-negative (DN; CD8^-^/CD4^-^), double-positive (DP; CD8^+^/CD4^+^), NK cells and Tγδ as follows: cells with non-null expression of *CD8A* and null expression of *CD4* were defined as CD8^+^ (and *vice-versa* for CD4^+^). Cells showing non-null expression of both genes were first classified as DP, then as doublets of CD4^+^ and CD8^+^ T cells as the average number of genes expressed per cell equaled close to the double of CD4^+^ or CD8^+^ T cell singlets. Due to notorious dropout events in single-cell data, cells lacking the expression of both markers were classified as follows: if a cell belongs to a cluster (taking a fine resolution of 10) in which the 75^th^ percentile expression of *CD8A* was higher than its 75^th^ percentile expression of *CD4*, it was classified as CD8^+^ (and *vice-versa* for CD4^+^). If the 75^th^ percentile expressions of both markers equal 0, the cells were classified as DN. Finally, cells with an average expression scores of all *TRG* and *TRD*-related genes higher than 0.5 (cutoff established after histogram visual inspection) were assigned to be Tγδ cells. We also characterized the T-cell repertoire of CD45^+^-sorted cells by scTCR-sequencing (VDJ) and compiled additional Tγδ cells for which TCR gamma or delta chains were found. In the general clustering of all CD45^+^-sorted cells, NK cells were clustering close to the CD8^+^ T cells and were annotated as NK1 and NK2 using the Zilionis *et al*. (*39*) centroid annotation by the *singleR* function from the *singleR* package.

The clustering of CD4^+^ T cells was obviously formed by three distinct clusters whose gene markers were indicating CD4 CXCL13 (T follicular-helper) cells (*CXCL13^+^*, *CD40LG^+^*, *BCL6^+^*, *CD200^+^*), Tregs (*FOXP3^+^*, *CTLA4^+^*, *IL2RA^+^*) and CD4 T helper 1 (Th1) cells (*IL7R^+^*, *SELL^+^*, *LEF1^+^*). For the CD8^+^ T cell subtyping, CD8^+^ T cells and NK cells were integrated by sample as explained above and by removing the TCR genes in order to prevent clustering based on clonotypes. We used several methods to annotate the clusters: (1) We first performed differential gene expression and differential regulon/TF activity (see below) analysis which were computed using the *FindAllMarkers* function with a 0.7 resolution. (2) We computed signature score (using the *AUCell* R package) for cytotoxicity (*CCL3*, *CCL4*, *CST7*, *GZMA*, *GZMB*, *IFNG*, *NKG7*, *PRF1*), exhaustion (*CTLA4*, *HAVCR2*, *LAG3*, *PDCD1*, *TIGIT*) and naiveness/memory (*CCR7*, *LEF1*, *SELL*, *TCF7*) taken from the Table S3 of Jerby-Arnon *et al*. list of genes (*48*). We extracted the gene list from Figure 7 of Andreatta *et al*. (*21*) as specific markers. We used the prediction from the annotation coming from public studies, namely Yost *et al*. (*18*), Guo *et al*. (*19*), Zhang *et al*. (*20*) and Oliveira *et al*. (*9*).

A cluster displaying elevated levels of *CCR7*, *LTB*, *SELL* and *IL7R* gene expression, high TCF7 and LEF1 regulon activity, high concordance with Oliveira *et al*. (*9*) predictions of naive cells was then named CD8 naive-like. Two clusters were displaying signs of stress or apoptosis: one with elevated levels of mitochondrial-related genes (gene names starting by *MT-*), expression of the *MALAT1* gene and lower number of genes expressed per cell was then named CD8 low-quality and another one with overexpression of heat-shock protein (HSP) genes and *DNAJ*-related genes which we named CD8 HSP. A cluster displaying concomitant expression of *CD8A* and *FOXP3* genes, without any signs of higher number of genes per cell (which could highlight doublets of CD8^+^ T cells and CD4 Tregs) and high concordance with Oliveira *et al*. (*9*) predictions of Treg-like cells was then named CD8 FOXP3. A cluster was obviously driven by the high expression of type-I interferon genes (*ISG15*, *MX1*, *IFI16*, *IFIT3*, *IFIT1*, *ISG20*, *OAS1*) and was then named CD8 ISG. A small cluster was found very close to the one of NK cells and displaying *CD8A* expression with co-expression of NK cells marker such as *KLRC2* and *KLRD1*, and high concordance with Oliveira *et al*. (*9*) predictions of NK-like cells was then named CD8 NK-like. Another small cluster that was captured at a finer resolution of 2, that displayed high expression of *CX3CR1* and *FGFBP2*, high levels of cytotoxic and low levels of exhaustion signatures, high concordance with Guo *et al*. (*19*) and Zhang *et al*. (*20*) predictions of CD8_CX3CR1 cells was then named CD8 CX3CR1. Four clusters with similar gene expression profiles, displaying high expression of *GZMK* (a marker for effector memory cells according to Andreatta *et al*. (*21*), modest cytotoxic and low exhaustion signature levels, high concordance with Oliveira *et al*. (*9*) predictions of CD8 Effector-Memory (EM) cells were then combined and named CD8 EM-like. Two clusters showing obvious signs of exhaustion with high levels of *HAVCR2* and *PDCD1* expression, high exhaustion signature levels, and elevated EOMES regulon activity were first classified as exhausted T cells then subclassified as follows: one cluster has higher *HAVCR2*, a typical sign of late exhaustion, and high concordance with Oliveira *et al*. (*9*) predictions of Terminal-Exhausted cells and was then named CD8 Tex; and the other cluster displayed higher levels of *TOX*, *XCL1*, *XCL2* and *CRTAM*, which are markers of precursor exhaustion according to Andreatta *et al*. (*21*), lower activity of TBX21 regulon and high concordance with Oliveira *et al*. (*9*) predictions of Progenitor-Exhausted cells and was thus named CD8 Pex. Finally, three clusters were driven by proliferation markers. Since proliferation is not a T-cell state in itself, we generated an average expression profile (centroid) for our CD8^+^ T cell subsets and predicted the T-cell state of proliferating CD8^+^ T cells by automated annotation using this centroid and the *singleR* R package. The full list of differentially expressed genes and TFs per cell types appears in Supplementary Table 4 and was computed using the *FindAllMarkers* function from the *Seurat* package. Pseudotime analysis was performed using *Monocle3* as inferred in the *SeuratWrappers* R library.

#### High-resolution cell type annotation of myeloid cells

Myeloid cell subtyping was achieved by integrating per sample as explained above. We used several methods to annotate the clusters (1) We first performed differential gene expression which was computed using the *FindAllMarkers* function with a resolution of 1. We used the prediction from the annotation coming from Zilionis *et al*. (*39*) and we used macrophage subtyping with specific genes as published by the group of Sohrab Shah (*41*). We first annotated the dendritic cell (DC) according to the predictions made using the Zilionis *et al*. (*39*) centroids. DC clusters were obviously corresponding to and then wo forced the clustered to be attributed to DC1, DC2, DC3, MonoDC and pDC calls. We also ensured that specific DC markers were expressed in the corresponding DC subsets (*CLEC9A* in DC1, *CD207* in DC2, *CCL22* in DC3, *CLEC10A* in MonoDC and *LILRA4* in pDC). The monocyte/macrophage component was annotated using the overlap between differential gene expression per cluster and specific genes extracted in Vázquez-García *et al* (*41*). We found clusters overexpressing the *S100A8*, *TREM2*, *CXCL9* and Complement genes, for which we then attributed the names Macro S100A8, Macro TREM2, Macro CXCL9 and Macro C1Q, respectively. A small population clustering close to the macrophages with high *FCGR3A* expression (the gene encoding CD16) was annotated as CD16 Monocytes. As for the CD8^+^ T cells, we also isolated clusters with mitochondrial gene expression and low number of genes expressed per cell which we attributed to be Macro low-quality, as well as a cluster overexpressing type-I interferon genes that we further named Macro ISG. We isolated a cluster with co-expression of both myeloid markers and T cell markers, and for which the number of genes was significantly higher than for the macrophage, that we attributed to Myeloid-T cell doublets. As for the CD8^+^ T cell, we generated myeloid cell specific average expression profiles and reannotated proliferating cells (cluster with high expression of cell cycle genes such as *MKI67*) using the *singleR* package. We did not find any mast cells, neutrophils or basophils in our data. The full list of differentially expressed genes and TFs per cell type appears in Table S4 and was computed using the *FindAllMarkers* function from the *Seurat* package.

#### M1 and M2 macrophage state classification

In order to attribute an M1 and/or M2 state to macrophages, we first computed gene signature scores for M1 and M2 taken from Azizi *et al.* (*8*) (ref). The cells were categorized into positive (M1^+^ and M2^+^) or negative (M1^-^ and M2^-^) for each signature if they were higher, respectively lower, than the median of each signature score for all macrophages. Macrophages were considered in M1 state if they were M1^+^/M2^-^ or in M2 state if they were M1^-^/M2^+^. Some cells were positive or negative for both categories.

#### High-resolution cell type annotation of B cells

B cell subtyping was achieved by integrating per sample as explained above. We used several methods to annotate the clusters. We first performed differential gene expression which was computed using the *FindAllMarkers* function with a resolution of 0.6. We got inspired by the B-cell subtyping performed in a pan-cancer single-cell study (*59*). We computed an immunoglobulin (Ig) signature score (using the *AUCell* R package) by capturing all genes whose names started with *IGH* or *IGL*. Four major subtypes of B cells were found in our dataset and could be annotated using specific markers: Plasma cell overexpressing *MZB1*, *JCHAIN*, *SDC1* and displaying high levels of the Ig signature. Naive B cells characterized by high levels of *FCER2*, *TCL1* and *IGHD*. Memory B cells were characterized by higher expression levels of *CD27* and less naive B-cell marker. We also found a germinal center (GC) population characterized by expression of *CD38* and *MEF2B*. Isolated small populations clustered separately from the Memory B cells but nevertheless expressing memory B-cell markers (*CD27*). These clusters overexpressed specific immunoglobulin chains, we thus thought they correspond to clonally expanded B cells but kept their annotation as Memory B cells. As for the myeloid subtyping, we found doublets of B cells with both myeloid and T cells exhibiting more expressed genes per cell. We also found proliferating B cells that were reannotated as described for myeloid and CD8^+^ T cells and low-quality B cells characterized by high expression of mitochondrial genes and lower number of expressed genes per cell. The full list of differentially expressed genes and TFs per cell type appears in Table S4 and was computed using the *FindAllMarkers* function from the *Seurat* package.

#### Annotation of the full sorted viable cell dataset using CD45^+^-sorted annotation

We first ensured about the high consistency between our baseline scRNAseq datasets by performing differential gene expression for major immune cell populations between Rs and NRs in both the CD45^+^-sorted (5’GEX) and total TME scRNAseq (3’GEX) datasets. Log fold-change correlations between the two scRNAseq libraries yielded highly similar DGE, thus allowing us to confidently utilize the CD45^+^-enriched data for the relative profiling and phenotyping of the immune compartment and the total TME data for the malignant phenotyping and quantification of cell stoichiometries. After the CD45^+^-sorted dataset (5’GEX) was annotated at higher resolution, we then generated averaged gene expression profiles (centroids) per cell subtype and used them to annotate the sorted viable cell dataset (3’GEX) using the *singleR* package. Briefly, we isolated major cell populations (CD4^+^ T cell, CD8^+^ T cell, myeloid and B cells) and performed automated annotation using *singleR* by only using centroids from the specific populations (i.e. CD4 Th1, CD4 CXCL13 and Tregs for the annotation of CD4^+^ T cells). The final annotation was not kept as predicted by the package but instead we went into fine resolution clusters and attributed each cluster the cell type whose prediction was the most abundant.

#### Doublet annotation

In order to deconvolute the cell subtypes composing the doublets, we created averaged expression profiles derived from singlets and, using the *singleR* tool that annotates cells based on correlation with a pre-existing gene expression profile, we guided the heterotypic doublet annotation into single cells of finer subtypes (i.e. a doublet involving a T cell and a myeloid cell was annotated into both a T cell and a myeloid state).

### Gene signature analysis

For single-cell data analysis and unless otherwise mentioned, gene signature scores were computed using the *AUCell* package. For bulk RNA sequencing data and bulked scRNAseq data, gene signature scores were computed using single-sample geneset enrichment anylsis (ssGSEA) method as inferred in the *gsva* function from the *GSVA* R package (with mx.diff=FALSE and ssgsea.norm=FALSE parameters). Individual gene signatures were taken from cited publications and collections were selected from the MSigDB portal (http://www.broadinstitute.org/gsea/msigdb); we used the Hallmarks and the Reactome collections. Differential analysis of gene signature scores was achieved using the regularized linear model as implemented in the *limma* package. Enrichment Barcode plot were performed using the *barcodeplot* function from the *limma* R package. The CD8^+^ T cell tumor-reactivity signature used for doublet characterization is derived from another study.

### Transcription factor activity in single-cell data

The transcription factor activity was estimated using the regulon signature of each transcription factor. Regulons were inferred using the *SCENIC* pipeline (https://scenic.aertslab.org) which integrates three algorithms (*grnBoost2*, *RcisTarget* and *AUCell*) corresponding to three consecutive steps:

Step 1: First, a gene regulatory network (GRN) was inferred from all tumors and ACT products transcriptomic together using grnBoost2, a faster implementation of the original Genie3 algorithm. grnBoost2 takes as the input scRNAseq transcriptomics data to infer causality from the expression levels of the transcription factors to the targets based on co-expression patterns. Basically, the prediction of the regulatory network between n given genes is split into n different regression problems and expression of a given target gene was predicted from the expression patterns of all the transcription factors using tree-based ensemble methods, Random Forests or Extra-Trees. The ranking of the relative importance of each transcription factor in the prediction of the target gene expression pattern is taken as an indication of a putative regulatory event. The aggregation targets into raw putative regulons was done using the runSCENIC_1_coexNetwork2modules function from the *SCENIC* R package with default parameters.

Step 2: Co-expression modules (raw putative regulons, i.e. sets of genes regulated by the same transcription factor) derived from the GRN generated in Step 1 are refined by removing indirect targets by motif discovery analysis using cisTarget algorithm and a cis-regulatory motif database. In particular, we used hg19-500bp-upstream-7species.mc9nr.feather and hg19-tss-centered-10kb-7species.mc9nr.feather databases. The motif database includes a score for each pair motif-gene, which allows the generation of a motif-gene ranking. A motif enrichment score is then calculated for the list of transcription factor selected targets by calculating the Area Under the recovery Curve (AUC) on the motif-gene ranking using the *RcisTarget* R package (https://github.com/aertslab/RcisTarget). If a motif is enriched among the list of transcription factor targets, a regulon is derived including the target genes with a high motif-gene score.

Step 3: Finally, *AUCell* was used to quantify the regulon activity in each individual cell (https://github.com/aertslab/AUCell). *AUCell* provides a AUC score for each regulon and cell; we discarded regulons with less than five constituent elements, as the estimation of the activity of small regulons is less reliable. Four the calculation of the AUC, the parameter *aucMaxRank* of the AUCell_calcAUC function was set with a fixed number of 1500 of features.

### Differential gene expression, regulon and pathway analyses

During the process of annotating the cells and to find markers of cell population like the one appearing in Table S4, we used the *FindAllMarkers* function from the *Seurat* package by using the following parameters: only.pos = TRUE, min.pct = 0.25 and logfc.threshold = 0.25. To derive gene expression, regulon activity and pathway score differential analyses according to clinical status, we performed linear regressions as inferred in the *lmFit* function of the *limma* R package. The mitochondrial and ribosomal contents were used as covariates in the regressions. Only the genes with average expression higher than a fixed threshold of 0.3 were kept in the final DEG table. No filtering was used for regulon and pathway analyses.

In differential analysis involving clinical status, we report the comparison of log fold-change analyses derived from single-cell based with those of patient-based data (by averaging the data). This approach helps to minimize the weight from variation of cell numbers among patients. This reassured us that our single cell-based observations were representative of all patients of each clinical response category and were not driven by outstanding cases. Statistical *p*-values for both single-cell and patient-based analyses appears in Table S2.

### Reactome pathway enrichment analysis

The 30 most upregulated or downregulated genes (computed by averaging logFC values from single-cell and patient-averaged analyses) according to clinical status were subjected to Reactome pathway enrichment analysis as follows: these genes were first converted into Entrez ID using *mapIds* from the *AnnotationDbi* package then were subjected to Reactome enrichment analysis using the *enrichPathway* function *ReactomePA* package. Reactome pathways with q-values lower than 0.01 were kept.

### Ligand-Receptor Interaction analysis

We developed our own methodology to infer cell-cell interaction considering cell type stoichiometry and expression of ligand and receptors in both cell types as described below. We used a database of ligands-receptors (LR) that was initially taken from the *SingleCellSignalR* package (LRdb.rda file). We isolated five different pathways from this database: the complement pathway that was isolated by capturing LR pairs containing the word ’Complement’ in the “ligand.name” or “receptor.name” columns. Similarly, for the Interferon and Interleukin pathways by using the words ’Interferon’ and ’Interleukin’, respectively. The co-stimulation and co-inhibitory gene list was extracted from Figure 1 of Chen *et al*. (*60*) and LR pairs containing these genes were isolated from the LR database. Similarly, Chemokine gene list was taken from Hornburg *et al*. (*61*) and LR pairs containing these genes were isolated from the LR database. To ensure capturing all existing interactions, the LR pairs for the five selected pathways were then enriched using another database (human_lr_pair.rds) taken from the *CellTalkDB* package. LR pairs with same receptor or ligand that were not present in the initial LR database was appended to it. Final list of LR pairs for the five selected pathways appears in the Table S6.

We computed cell-cell interaction scores based on the average expression of a ligand and its cognate receptors in two specific cell type (finer resolution) weighted by their corresponding proportions (real stoichiometry extracted from the sorted viable cell dataset (3’GEX)) and computed every possible combination of ligands and receptors in all cell types as follows:

*Interaction score* = (*Prop cellType*1)(*AvgExpGene*1 *in CelType*1)(*Prop CellType*2)(*AvgExpGene*2 *in CelType*2)

`Where Prop is the proportion of this cell type in the full sorted viable cell dataset and AvgExp is the average expression of the gene in this particular cell type`

We then run Student’s t-test according to clinical status for all these possible combinations and extracted significantly different interaction scores between responders and non-responders (uncorrected *p*-value <= 0.05). Significant interactions were then selected for plotting using heatmap and circos plotting. These analyses were run in the sorted viable cell dataset for baseline tumor and in the CD45^+^-sorted dataset for post-ACT T30 data as sorted viable data was not available for this time-point.

### TCR/VDJ related analyses

5’ scTCRseq (VDJ) data were used to compute richness and clonality metrics in CD8^+^ T cell subsets, as well as to track the TCR chains across several CD8^+^ T cell subsets. Only TCR-β chains (The concatenation of the two TCR-β chains) was considered for the computations. The richness per CD8^+^ T cell subsets was considered as the number of unique TCR-β chains per subset normalized by the number of cells in this CD8^+^ T cell subset. The clonality was computed as 1-Pielou’s index (*62*). We also plotted the number of unique TCR-β chains shared between pairs of CD8^+^ T cell subsets.

### Plotting description and statistical analyses

All heatmaps were performed using the *pheatmap* function from the *pheatmap* R package. Violin plots were achieved by first plotting the violins using the *violin_plot* function from the *plotrix* R package, followed by addition of symbols by using the *stripchart* function from the *graphics* R package. Plotting of UMAP and Dot plot shoing expression of genes per cell subtypes were were achieved using the *DimPlot*, respectively *DotPlot* function from the *Seurat* R package. UMAP highlighting gene expression or signature score in density were done using the *plot_density* function of the *Nebulosa* R package. UMAP showing performance of two signature scores at the same time was plotted using the *plot* function from the *base* R package and the script to convert the two gene signatures scores for exhaustion and cytotoxicity into a double-color code was taken from Wu *et al*. (*57*). Scatter plots, Ridge plots and Fraction plots were performed using the *ggplot2* R package. Alluvial plots were performed using the *alluvial* R package. Bar plots and line plots tracking points at different time-points were done using the default *graphics* R package. Circos plots were performed in R using the *circlize* package. Figures were reprocessed using Adobe Illustrator 2020 for esthetical purposes. The method used for statistical analysis appears directly in the method section related to the specific technique that was used or directly in the figure legend. ROC analyses were performed using the *prediction* function from the *ROCR* package. The true-positive rate, false-positive rate and area under the curve (AUC) were then extracted using the *performance* function of the *ROCR* package. Statistical tests used for specific analyses directly appear in the corresponding figure legends.

---

**Figure S1.**
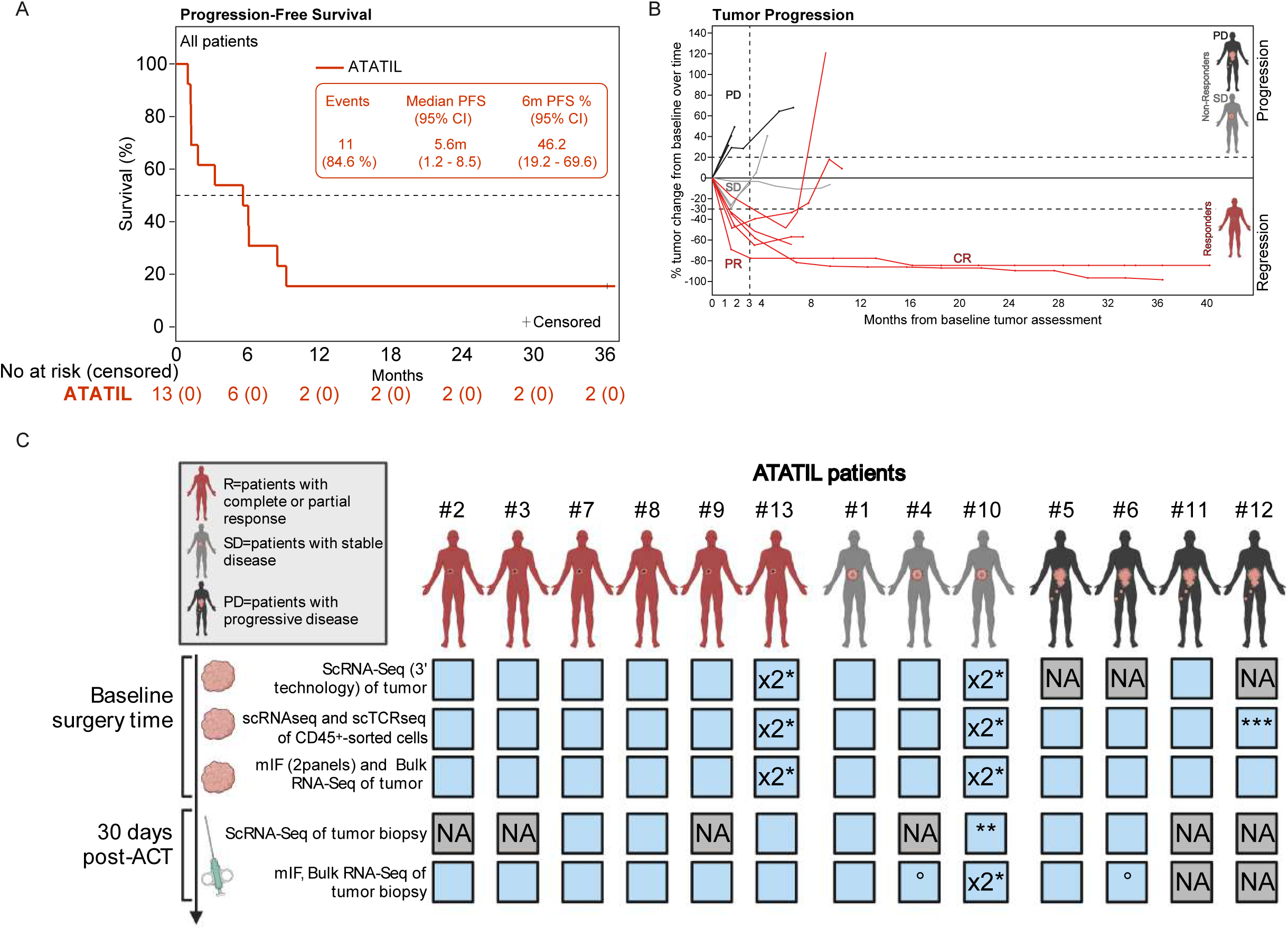
Sample availability and clinical features of the ATATIL cohort. (Related to Figure 1). (A) Progression-free survival (PFS) for the ATATIL cohort by reverse Kaplan-Meyer method. Spider plot showing percentage of tumor change relative to baseline over. At day 30 (first tumor assessment scan) two patients had stable disease (SD) based on RECIST 1.1, however, at the next scan a progressive disease (PD) was recorded. So, for these two patients, PD is considered as the best response. (**C**) Available material is depicted in light blue while unavailable material appears in gray. (*) Two baseline tumors were surgically removed. Samples were analyzed separately for bulk and pooled for single-cell analyses. (**) Two tumor biopsies were obtained 30 days post-ACT: one responding and one lesion in progression. For bulk analyses, both lesions were available whereas for single-cell, only the responding lesion could be processed. (***) scRNAseq (5’ technology) was done on sorted viable cells rather than CD45^+^-sorted cells due to high infiltration. (°) mIF data non-available.

**Figure S2.**
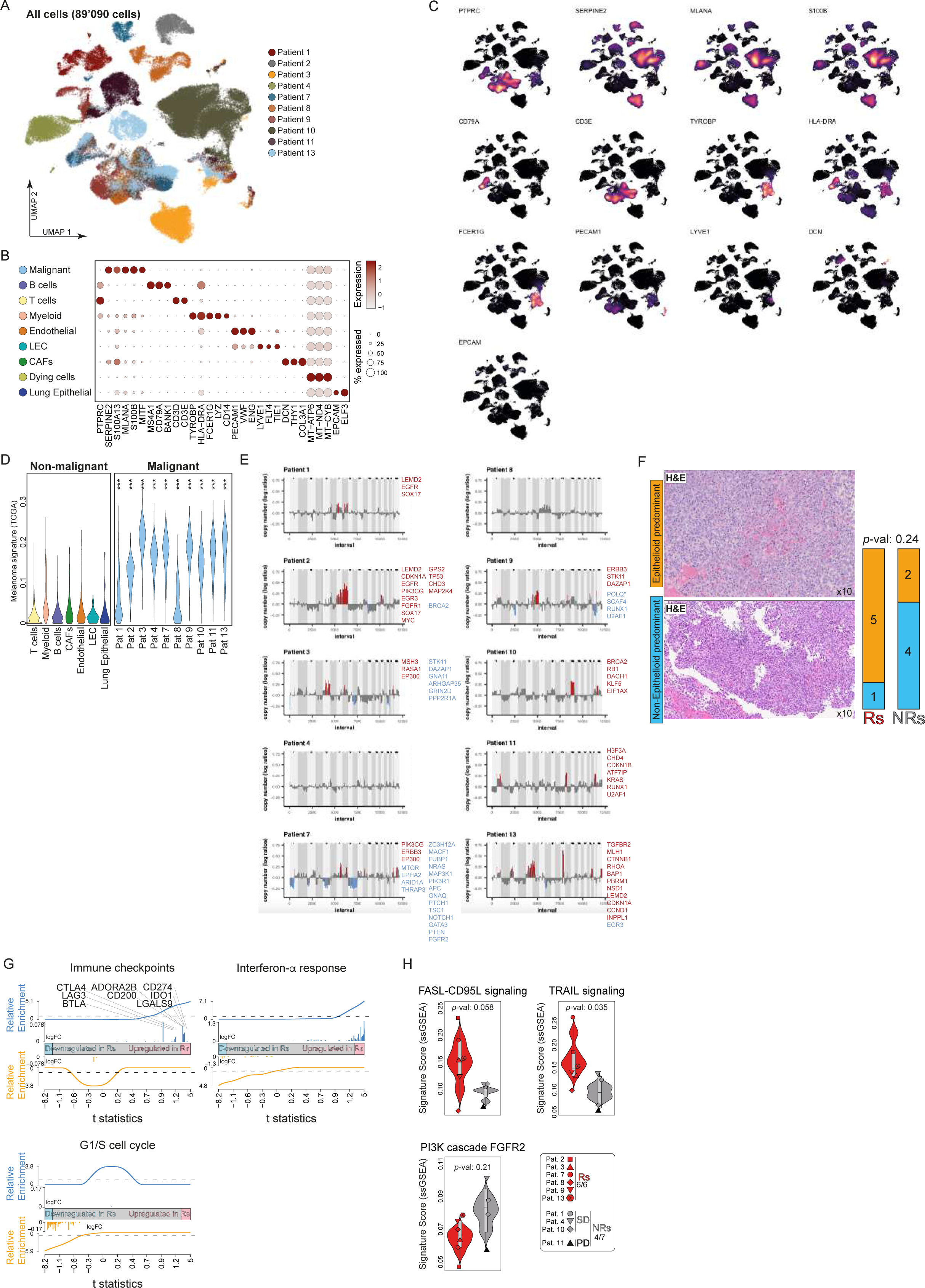
Characterization of the malignant compartment. (Related to Figure 2). (**A**) UMAP projection of sorted viable cell scRNAseq data (3’GEX) from ten baseline melanoma highlighting the patient identity. (**B**) Dot plot showing main lineage gene expression according to the main cell types found in sorted viable single-cell data. (**C**) UMAP density plot showing the smoothed expression of main lineage gene markers. (**D**) Scoring of a melanoma signature derived from TCGA analysis in non-malignant cell types and in expectedly-malignant private clusters of each patient. Statistical analysis performed by ANOVA followed but *post-hoc* Tukey test for multiple comparison correction. (***) indicates corrected *p*-values lower than 0.001. (**E**) CNV inference from gene expression showing genomic amplification (in red) and deletion (in blue). Oncogenic or tumor suppressor driver genes falling in these regions are highlighted. (**F**) Example of hematoxylin and eosin (H&E) staining showing epithelioid and non-epithelioid tissues and their quantification according to clinical status. Statistics performed by Fisher’s Exact testing. (**G**) Barcode plot displaying where genes involved in selected pathways rank in the list of differential gene expression between malignant cells of Rs and NRs. (**H**) Violin plots showing signature scores in malignant cells averaged per patient for the indicated Reactome pathway according to clinical response. Statistical significance was extracted from analysis in Figure 2E.

**Figure S3.**
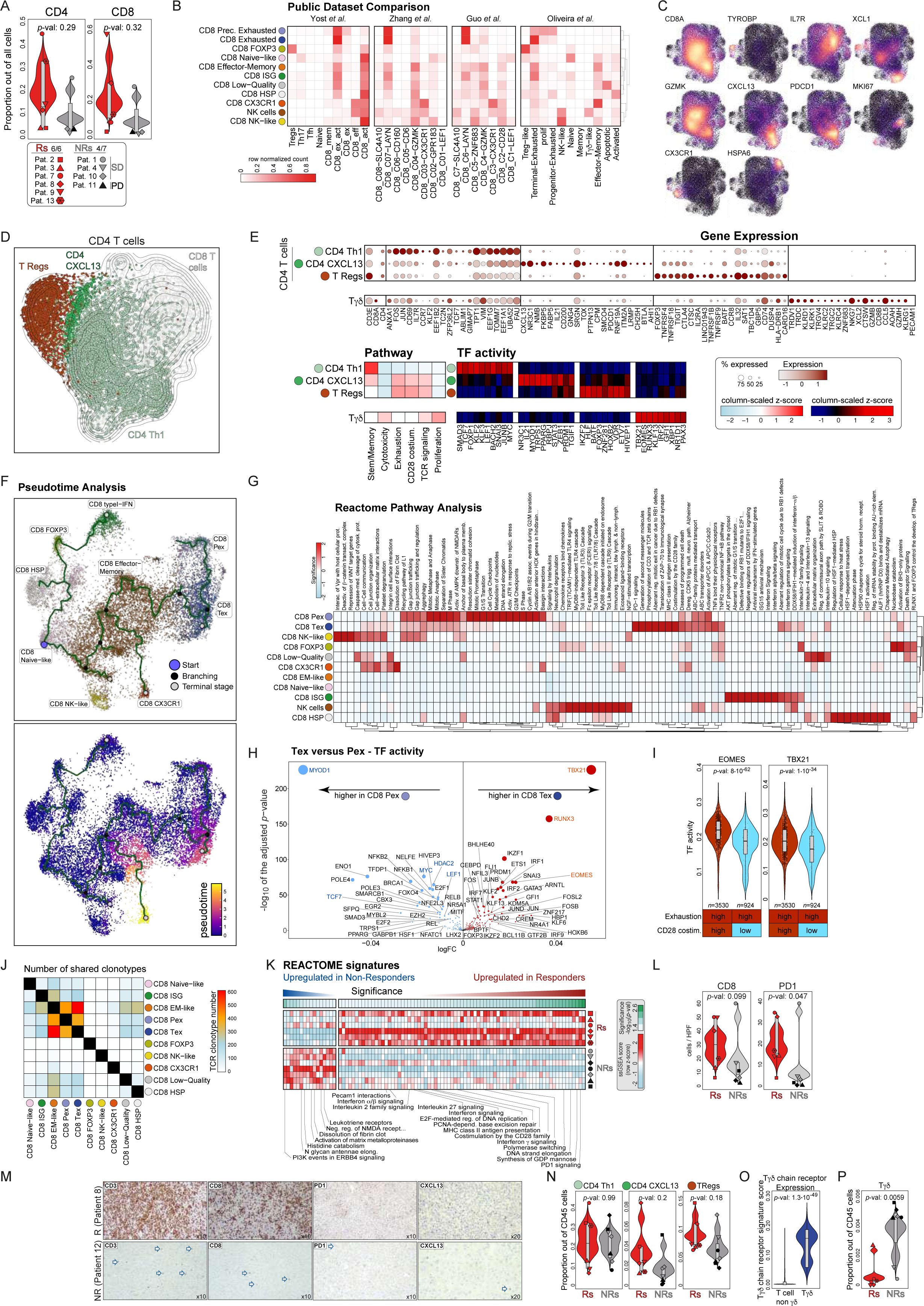
Characterization of the T-cell compartment. (Related to Figure 3). (**A**) Violin plots of CD4 T and CD8 T cell proportions over all cells in sorted viable cell data according to clinical response. Statistical significance was assessed using Student’s t-test comparing Rs versus NRs. (**B**) Comparison of ATATIL CD8^+^ T cell annotation with the profiles published by other studies. (**C**) UMAP density plot showing the smoothed expression of CD8^+^ T cell-related gene markers in integrated CD8^+^ T cell/NK clustering of CD45^+^-cell sorted data. (**D**) UMAP projection showing sub-clustering of T and NK cells from sorted viable cell scRNAseq data (3’GEX) from 10 baseline tumors. CD4^+^ T cell subsets are highlighted. (**E**) Characterization of the CD4^+^ T and Tγδ cell populations: at the top, dot plot showing discriminant gene expression markers. At the bottom left, gene signature scores for stemness/memory, cytotoxic and exhaustion (taken from Jerby-Arnon *et al*.), TCR signaling and costimulation mediated by CD28 (Reactome) and proliferation per T cell subtype appear. At the bottom right, heatmap showing the most discriminant transcription factors for each population. (**F**) Pseudotime analysis of the CD8^+^ T cell populations performed by *Monocle3*. Top plot showing the UMAP projection and the trajectory starting from CD8 naïve-like cells. Branching and terminal stages are indicated as shown in the figure. Bottom plot showing the UMAP colored by the pseudotime values of each cell. (**G**) Heatmap showing clustering of the most enriched Reactome pathways per CD8^+^ T cell subsets for each CD8^+^ T cell subset using the CD45^+^-cell sorted data. (**H**) Volcano plot displaying differential transcription factor/regulon analysis between CD8 Tex and CD8 Pex. (**I**) Regulon activity for EOMES and TBX21 in CD8^+^ T cells that were categorized into high and low for exhaustion signature (Jerby-Arnon *et al.*) and CD28 costimulation (Reactome) using median cutoff. Statistical significance was assessed using Student’s t-test comparing Rs versus NRs. (**J**) Number of TCR clonotypes (TCR β-chains that were found in two different states (at least in one cell of each cell subset). (**K**) Reactome pathway analysis showing selected pathways that are the most differentially represented between CD8^+^ T cells from Rs versus NRs. Statistical significance was assessed using linear modeling as described in the methods. (**L**) Violin plot showing quantification of CD8^+^ and PD1^+^ (from Figure 3J) cell densities (cell count/HPF) between Rs and NRs. Statistics performed using one-sided wilcoxon tests. (**M**) Example of immunohistochemistry (IHC) staining showing CD3^+^, CD8^+^ and PD1^+^ cells at 10x magnification and CXCL13^+^ cells at 20x magnification on pre-REP tumor material by Pathology evaluation. (**N**) Violin plots of CD4^+^ T cell subtype proportions over CD45^+^ cells in CD45^+^-sorted data according to clinical response. Statistical significance was assessed using Student’s t-test comparing Rs versus NRs. (**O**) Violin plot showing that Tγδ cells display a significantly higher score for *TRD* and *TRG*-related genes compared to non-Tγδ cells. Statistical significance was assessed using Student’s t-test comparing Rs versus NRs. (**P**) Violin plots of Tγδ cell proportions over CD45^+^ cells in CD45^+^-sorted data according to clinical response. Statistical significance was assessed using Student’s t-test comparing Rs versus NRs.

**Figure S4.**
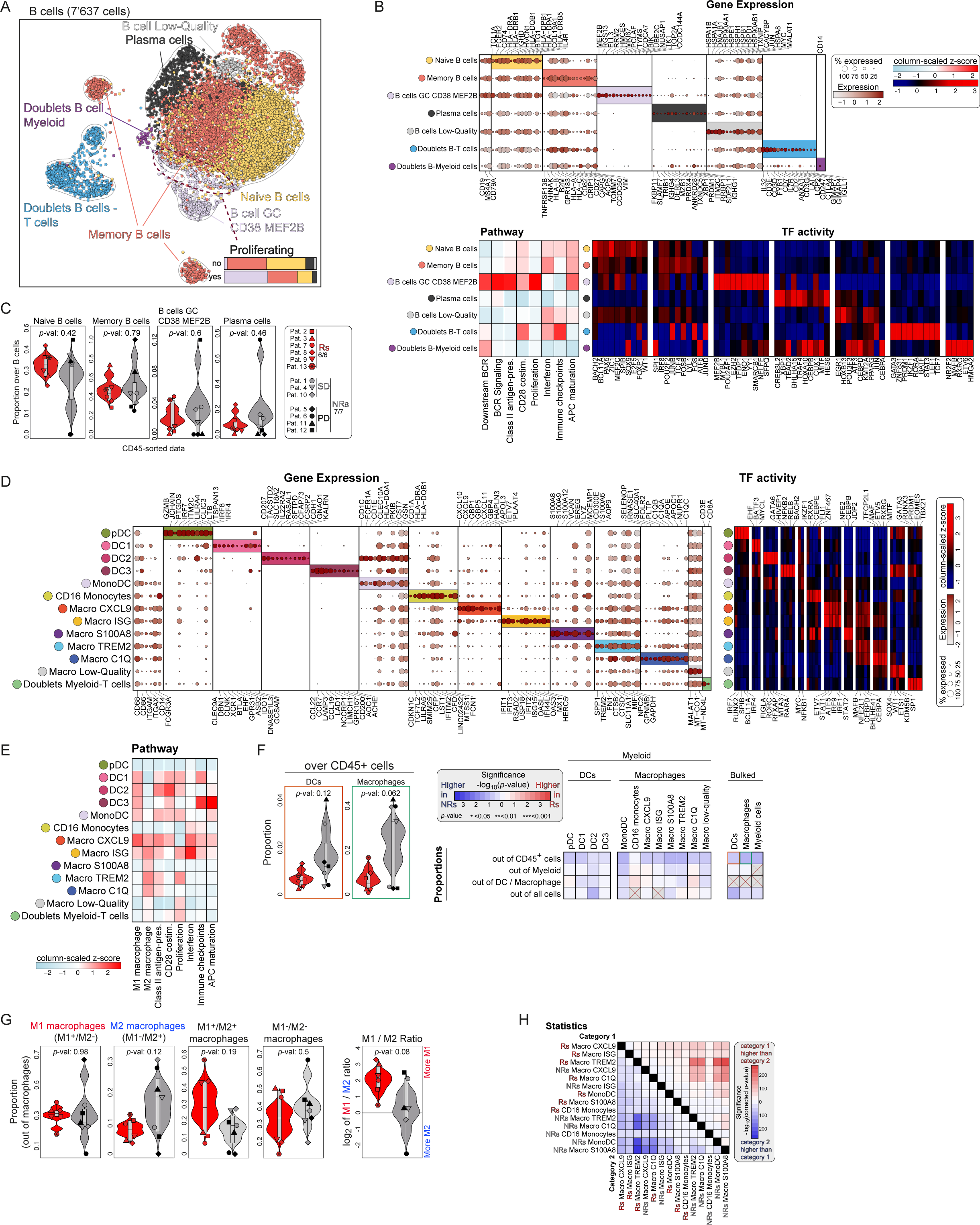
Characterization of the myeloid and B-cell compartments. (Related to Figure 4). (**A**) UMAP projection showing sub-clustering of B cells from CD45^+^-cell sorted data from thirteen baseline and seven post-ACT tumors. (**B**) Characterization of the B-cell populations: at the top, dot plot showing discriminant gene expression markers. At the bottom left, gene signature scores for downstream BCR signaling and BCR signaling (Reactome), class-II antigen-presentation (Reactome), costimulation mediated by CD28 (Reactome), Interferon (Reactome), Immune checkpoints and APC maturation (curated) and proliferation per B-cell subtype appear. At the bottom right, heatmap showing the most discriminant transcription factors for each population. (**C**) Violin plots of B-cell subtype proportions over total B cells in CD45^+^-cell sorted data according to clinical response. Statistical significance was assessed using Student’s t-test comparing Rs versus NRs. (**D**) Characterization of the myeloid cell populations: on the left, dot plot showing discriminant gene expression markers. On the right: heatmap showing the most discriminant transcription factors for each population. (**E**) Gene signature scores for M1/M2 macrophage polarization (Azizi *et al.*), class II antigen-presentation (Reactome), costimulation mediated by CD28 (Reactome), Interferon (Reactome), Immune checkpoints and APC maturation (curated) and proliferation per myeloid cell subtype. (**F**) Left: Proportion of bulked DC and macrophage populations out of CD45^+^ cells. Statistical significance was assessed using Student’s t-test comparing Rs versus NRs. Right: Heatmap showing the differences in the proportions of myeloid subsets according to clinical response and out of the respective set of cells as indicated in the figure. The heatmap display directed *p*-values (in -log_10_) with red colors indicating higher proportions in Rs and blue colors indicating higher proportions in NRs (using Student’s t-test). Asterisks denote significance as highlighted in the legend. (**G**) Left: proportions of M1 and/or M2 macrophages out of macrophages and according to clinical response. Right: Ratio between M1 and M2 macrophages. Statistical significance was assessed using Student’s t-test comparing Rs versus NRs. (**H**) Statistical analysis of macrophage clinical response signature score per macrophage subsets split according to response (related to Figure 4E). Statistics performed using Student’s t-test, adjusted using Bonferroni correction and plotted as a heatmap.

**Figure S5.**
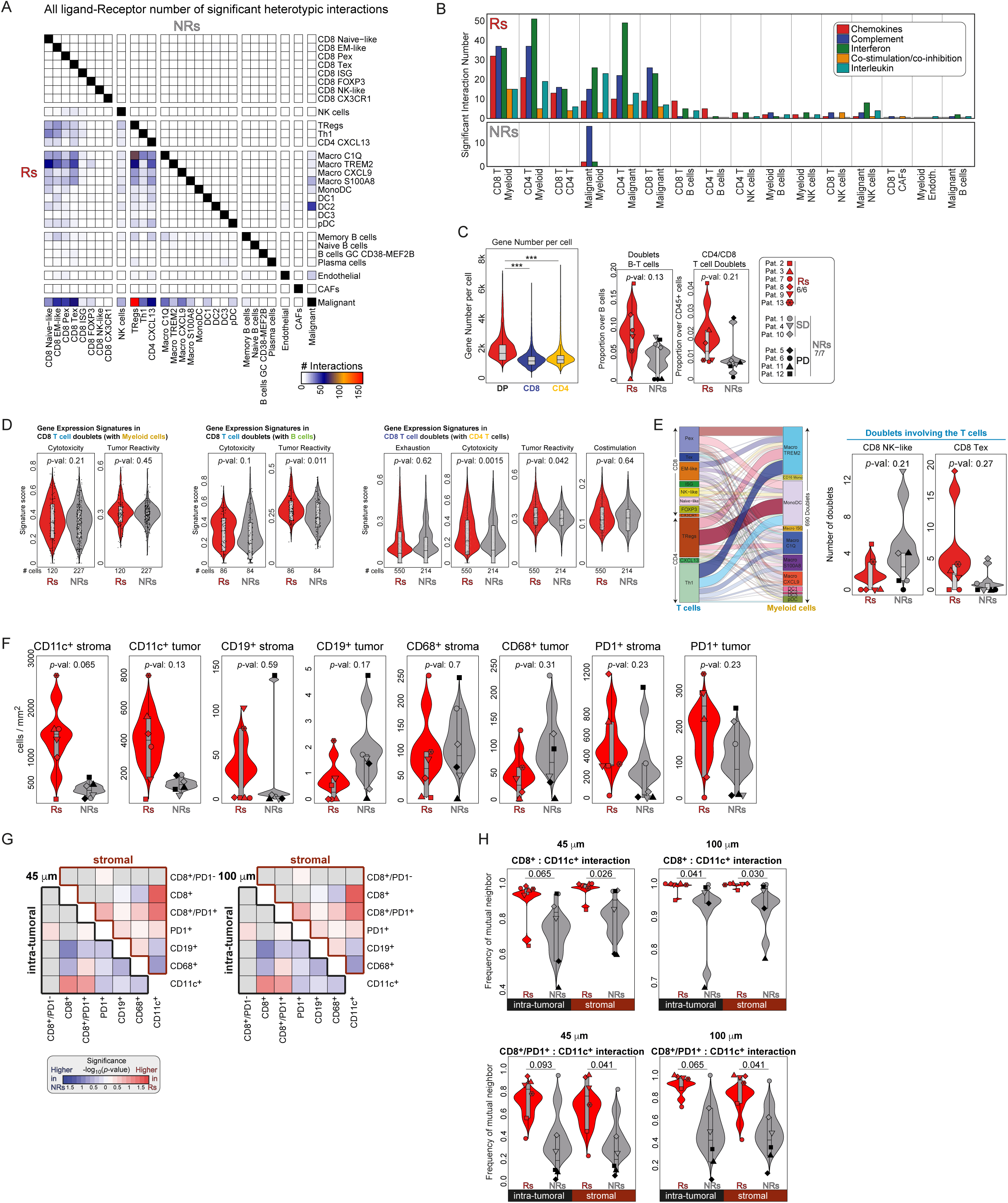
Characterization of the TME crosstalk. (Related to Figure 5). (A) Heatmap displaying the number of significant ligand-receptor (from the entire LR database) interactions between cell subsets in responders (lower-left part of the heatmap) and non-responders (higher-right part of the heatmap). (**B**) Number of significant (unadjusted *p*-value < 0.05) ligand-receptor pair interaction according to the indicated main cell types categorized into five different pathways and split by clinical response. (**C**) Left: number of genes expressed per cell in CD4^+^ T cell:CD8^+^ T cell doublets, and CD4^+^ T and CD8^+^ T singlets. Statistical significance was assessed using Student’s t-test comparing Rs versus NRs. *** denotes *p*-values lower than 0.001. Right: violin plots showing the proportions of B cell:T cell doublets and CD4^+^ T cell:CD8^+^ T cell doublets in CD45^+^-cell sorted data. Statistical significance was assessed using Student’s t-test comparing Rs versus NRs. (**D**) Selected signature scores computed in doublets involving CD8^+^ T cells and myeloid cells (left), B cells (middle) and CD4^+^ T cells (right). Statistical significance was assessed using Student’s t-test comparing Rs versus NRs. (**E**) Left, alluvial plot showing cell subset deconvolution in the myeloid cell-T cell doublets. Right, examples of specific cell subsets involved in the doublet deconvolution according to clinical response. Statistical significance was assessed using Student’s t-test comparing Rs versus NRs. (**F**) Violin plot of cell density (cell/mm^2^) between Rs and NRs. Statistics performed using wilcoxon tests. (**G)** Heatmaps showing the differences in the frequency of mutual neighbors between cell types at different neighboring radii. The heatmaps display directed *p*-values (in -log_10_ and using Student’s t-test) for pairwise comparisons with red colors indicating higher frequencies in Rs and blue colors indicating higher frequencies in NRs. (**H)** Violin plots displaying frequency of mutual neighbors at different neighboring radii between the indicated cell types, according to clinical response and split by stromal or intratumoral areas. Statistical significance was assessed using Student’s t-test comparing Rs versus NRs.

**Figure S6.**
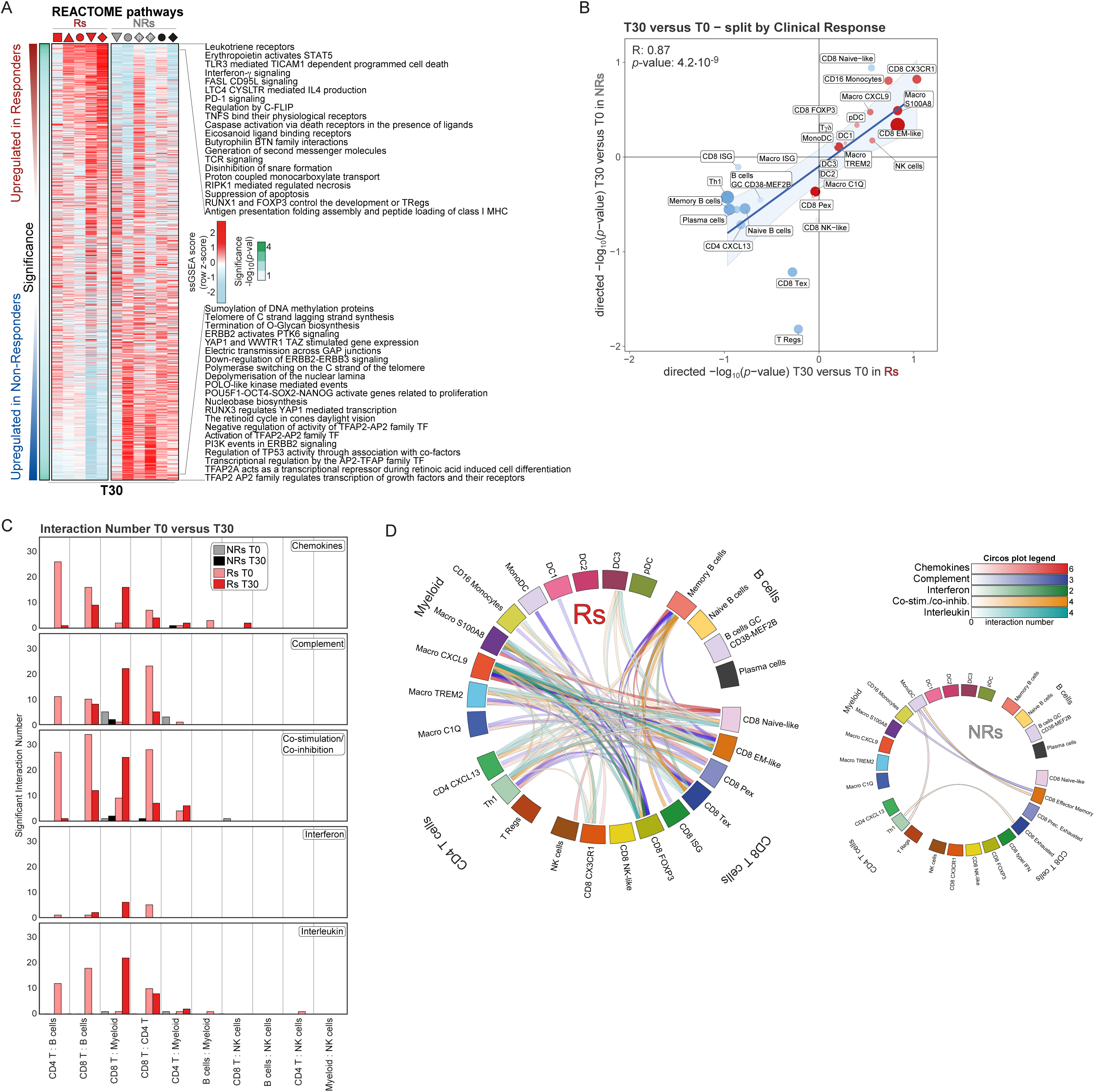
Characterization of the TME characteristics and progression post TIL-ACT (Related to Figure 6). (**A**) Reactome pathway analysis in bulk RNAseq of post-ACT T30 tumors showing most differentially represented pathways between Rs versus NRs. Statistical significance was assessed using linear modeling as described in the methods. (**B**) Differential analysis of cell subtype proportions between T30 and T0 and split by clinical response (directed -log_10_ of the *p*-value in responders on the x-axis and in non-responders on the y-axis) in CD45^+^-cell sorted single-cell data. (**C**) Number of significant (unadjusted *p*-value < 0.05) ligand-receptor pair interaction according to the indicated main cell types categorized into five different pathways and split by clinical response and by time when the biopsies were taken (T0 and T30). (**D**) Circos plots showing the finer cell subtypes involved in significant interactions (link) and the number of significant interactions (in color intensity) for the ligand-receptor pairs of five different pathways and split by clinical response for post-ACT T30 data.

**Table S1. (Supplementary_Table_1.xlsx). Clinical characteristics and survival analysis of patients enrolled in the ATATIL trial.**

**Sheet 1**: Patients are listed in columns and clinical characteristics referring to screening time-point in lines. ACT: adoptive cell-therapy; PS: performance status according to Eastern Cooperative Oncology Group (ECOG); LDH: lactate dehydrogenase level in blood (UI/ml). **Sheet 2**: Overall survival (OS) and progression-free survival (PFS) analysis according to age, gender, *BRAF* status, number of IL-2 doses received, LDH levels and neutrophil-lymphocyte ratio. Statistics were performed using univariate and multivariate Cox proportional hazard models. *P* stands for *p*-value and HR for hazard ratio.

**Table S2. (Supplementary_Table_2.xlsx). Differential Expressed Genes (DEG) per cell population according to clinical response**. Each sheet displays the DEG for each main cell population (malignant cells, CD8^+^ T cells, CD4^+^ T cells, Macrophages, DCs and B cells). Statistical significance was assessed using linear modeling as described in the methods. Positive logFC indicates expression values that are higher in responders and *vice-versa*. The “_sc” and “_patient” labels indicate values for which the analyses were performed at single-cell and patient-averaged levels, respectively. When applicable, a “Clinical_Response_Signature_Selection” field indicates whether genes are included in the clinical response gene signature of the corresponding cell type.

**Table S3. (Supplementary_Table_3.xlsx). Differential Reactome pathway analysis according to clinical response**. (sheets 1 to 3): All Reactome pathway scores were computed for every cell and a differential analysis of these scores was performed according to response. Each sheet displays the differential analysis for the following main cell population: malignant cells, CD8^+^ T cells, CD4^+^ T cells. Statistical significance was assessed using linear modeling as described in the methods. Positive logFC indicates signature scores that are higher in responders and *vice-versa*. (sheets 4 to 6): Reactome Term enrichment analysis for the 30 most upregulated and downregulated genes in differential gene expression analysis for the following main cell populations: macrophages, DCs and B cells.

**Table S4. (Supplementary_Table_4.xlsx). Differential Expressed Genes (DEG) and TF (DE_TF) according to cell subsets in CD45^+^-cells sorted single-cell data.** List of differentially expressed genes per cell subtype performed using the *FindAllMarkers* function from the *Seurat* R package and filtering only positive genes with a log fold-change higher than 0.25. Each sheet displays the differential analysis performed within the indicated cell type or considering all TME cells together (“All_TME”).

**Table S5. (Supplementary_Table_5.xlsx). Baseline tumor (T0) Ligand-Receptor (LR) differential interaction analysis according to clinical response in the sorted viable single-cell data.** Manual LR interaction analysis showing significant (*p*-value <= 0.05 uncorrected Student’s t-test) interaction scores (see methods) between Rs and NRs for each LR pair in the indicated cell subsets. All possible LR pairs were tested.

**Table S6. (Supplementary_Table_6.xlsx). List of LR pairs for five different pathways involved in immune cell interactions.** The following selected pathways appear in separated sheets: Chemokines, Complement, Interferon, Costimulation/Coinhibition and Interleukin.

**Table S7. (Supplementary_Table_7.xlsx). Baseline tumor (T0) Ligand-Receptor (LR) differential interaction analysis according to clinical response in the sorted-viable single-cell data in five selected pathways.** Manual LR interaction analysis showing all LR interaction score (see methods) between Rs and NRs in the indicated cell subsets. Tested LR pairs were taken from five different pathways involving immune cell interactions (see Table S6).

**Table S8. (Supplementary_Table_8.xlsx). Differential Reactome pathways analysis according to clinical response in post-ACT biopsies.**

**Sheet 1**: All Reactome pathways scores were computed in bulk RNA sequencing data. Paired differential analysis was performed between T30 and T0 in Rs and NRs separately. LogFC and *p*-values are reported for all pathways. Positive logFC indicate signature scores that are higher in T30 biopsies. This table is related to Figure 6A. **Sheet 2**: All Reactome pathways scores were computed in bulk RNA sequencing data. Differential analysis was performed between T30 biopsies from Rs and T30 biopsies from NRs. Positive logFC indicate signature scores that are higher in T30 biopsies from responders. This table is related to Figure S6A.

**Table S9. (Supplementary_Table_9.xlsx). Day 30 post-ACT (T30) Ligand-Receptor (LR) differential interaction analysis according to clinical response in the sorted-viable single-cell data.** Manual LR interaction analysis showing significant (*p*-value <= 0.05 uncorrected Student’s t-test) interaction score (see methods) between Rs and NRs for each LR pair in the indicated cell subsets. Tested LR pairs were taken from five different pathways involving immune cell interaction (see Table S6). Each sheet displays the significant interaction found between major cell-types.

